# AUTS2 controls neuronal lineage choice through a novel PRC1-independent complex and BMP inhibition

**DOI:** 10.1101/2021.06.29.450402

**Authors:** Zhuangzhuang Geng, Qiang Wang, Weili Miao, Trevor Wolf, Jessenia Chavez, Emily Giddings, Ryan Hobbs, David J. DeGraff, Yinsheng Wang, James Stafford, Zhonghua Gao

## Abstract

Despite a prominent risk factor for Neurodevelopmental disorders (NDD), it remains unclear how *Autism Susceptibility Candidate 2* (*AUTS2*) controls the neurodevelopmental program. Our studies investigated the role of AUTS2 in neuronal differentiation and discovered that AUTS2, together with WDR68 and SKI, forms a novel protein complex (AWS) specifically in neuronal progenitors and promotes neuronal differentiation through inhibiting BMP signaling. Genomic and biochemical analyses demonstrated that the AWS complex achieves this effect by recruiting the CUL4 E3 ubiquitin ligase complex to mediate poly-ubiquitination and subsequent proteasomal degradation of phosphorylated SMAD1/5/9. Furthermore, using primary cortical neurons, we observed aberrant BMP signaling and dysregulated expression of neuronal genes upon manipulating the AWS complex, indicating that the AWS-CUL4-BMP axis plays a role in regulating neuronal lineage specification *in vivo*. Thus, our findings uncover a sophisticated cellular signaling network mobilized by a prominent NDD risk factor, presenting multiple potential therapeutic targets for NDD.

## Introduction

Neurodevelopmental disorders (NDD) result from compromised development of the central nervous system and encompass but are not limited to Autism Spectrum Disorders (ASD), Developmental Delay (DD), and Intellectual Disabilities (ID). *Autism Susceptibility Candidate 2* (*AUTS2*) has been identified as one of the genes that are most frequently disrupted by balanced chromosome abnormalities (BCAs) in neurodevelopmental disorders (Talkowski et al., 2012). Although genomic aberrations of *AUTS2* were first identified in zygotic twins with autism, it has been repeatedly found in patients diagnosed with various NDD (Beunders et al., 2016; Oksenberg and Ahituv, 2013). Disruption of *AUTS2* leads to common neurodevelopmental abnormalities, including microcephaly, developmental delay, and varying degrees of intellectual disability (Oksenberg and Ahituv, 2013). The importance of *AUTS2* in the developing central nervous system (CNS) has been studied in animal models. In zebrafish, the deletion of *auts2* has been shown to affect neurogenesis and craniofacial development (Beunders et al., 2013; Oksenberg et al., 2013). Mouse models further reveal the relationship between AUTS2 loss of function and subsequent defects in CNS development and aberrant behaviors (Gao et al., 2014; Hori et al., 2015; Hori et al., 2014). Furthermore, using mouse and human embryonic stem cells (ESC), it has been shown that deleting *Auts2* leads to differentiation defects in neuronal lineage *in vitro* (Monderer-Rothkoff et al., 2021; Russo et al., 2018). However, it remains unclear how AUTS2 regulates neuronal differentiation at the molecular level.

Expression of AUTS2 in the CNS peaks during early embryogenesis and declines after birth (Bedogni et al., 2010; Gao et al., 2014). AUTS2 exists predominantly in the nucleus (Bedogni et al., 2010; Gao et al., 2014), but one study suggests that its functions depend on its cytosolic expression (Hori et al., 2014). Our previous studies have shown that nuclear AUTS2 comprises a type 1 Polycomb repressive complex (PRC1-AUTS2) (Gao et al., 2012) and converts PRC1 from a transcriptional repressor to an activator (Gao et al., 2014). However, cytosolic AUTS2 seems essential for neuronal migration and neuritogenesis in the mouse cortex through the activation of Rac1 (Hori et al., 2014). To date, two major isoforms of AUTS2 have been identified, but their functional distinctions are not well understood (Monderer-Rothkoff et al., 2021). Additional factors that interact with AUTS2 also have been identified (Gao et al., 2014; Monderer-Rothkoff et al., 2021), but it is unclear how these mechanistically contribute to the AUTS2-mediated control of neuronal cell fate.

It has been shown recently that deletion of Pcgf5, which is the core component in the PRC1-AUTS2 complex, causes abnormal TGF-β signaling activation and a defect in mouse ESC neuronal differentiation (Yao et al., 2018). However, whether or not AUTS2 directly regulates TGF-β/BMP pathways during neuronal differentiation remains unknown. TGF-β signaling plays essential roles in cell fate determination (Massague, 2012). The differentiation of ESCs into neuroectodermal lineage requires the absence of TGF-β and its family members, bone morphogenetic proteins (BMPs), while mesoderm and endoderm differentiation rely on these signaling pathways (Watabe and Miyazono, 2009). Furthermore, the inhibition of TGF-β and BMP signaling greatly enhances the efficiency of neuronal differentiation from ESCs *in vitro* (Chambers et al., 2009). The activation of the TGF-β/BMP pathway leads to the phosphorylation of regulatory SMAD (R-SMAD) proteins, among which SAMD1/5/9 responds to BMP and SMAD2/3 to TGF-β. The phosphorylated R-SMADs translocate to the nucleus in association with SMAD4 (co-SMAD) and elicit specific transcriptional effects (Massague, 2012).

In the present study, we discovered that the long but not the short isoform of AUTS2 interacts with PRC1 components. Using CRISPR/Cas9-mediated gene editing, we generated mouse ESCs that lacked either long, short, or both isoforms of Auts2 and demonstrated that complete silencing of both Auts2 forms leads to dramatic defects in neuronal differentiation. Interestingly, this differentiation defect in Auts2-deficient cells is accompanied by an up-regulation of BMP signaling, a critical pathway involved in cellular differentiation. Mechanistically, with quantitative mass spectrometry analysis, we identified a novel protein complex comprised of AUTS2, WDR68, and SKI, which is specifically formed in neuronal progenitors and mediates the inhibition of BMP signaling during differentiation. The impact of *Auts2* on neuronal gene transcription and BMP signaling was recapitulated in cortical neurons with *Auts2* deletion. Further biochemical characterization revealed the involvement of the CUL4 E3 complex as the mediator, which promotes proteasomal degradation of BMP-specific regulatory SMAD1/5/9, thereby restricting the activation of BMP signaling during neuronal differentiation.

## Results

### 1) Auts2 is required for neuronal differentiation in mouse embryonic stem cells

To explore how Auts2 regulates neuronal differentiation, we used an *in vitro* ESC neuronal differentiation model as previously described (Bibel et al., 2004). E14 cells were used to form embryoid bodies (EB), which were then induced by retinoic acid to differentiate into neuronal progenitor cells (NPC). The success of our differentiation was evidenced by the reduction of Oct4 and Nanog protein levels, two pluripotent markers, and the increase of Pax6 and Nestin, two NPC markers, during differentiation (Fig. 1a). Interestingly, even though protein levels remained constant in most of the tested PRC1-AUTS2 complex components, including Pcgf3, Pcgf5, Ring1b, and Wdr68, both the long and short isoforms of Auts2 (Auts2-L and Auts2-S) were increased in NPCs (Fig. 1a). These Auts2 isoforms have been previously observed by others (Beunders et al., 2013; Hori et al., 2014). *Auts2-S* is derived from an alternative transcription start site and shares the same reading frame with *Auts2-L* (Fig. S1a) (Beunders et al., 2013). Through immunoprecipitation (IP) experiments in HEK293 cells, we discovered that, unlike AUTS2-L, AUTS2-S does not associate with RING1B and PCGF5, the core components of the PRC1-AUTS2 complex (Fig. 1b). However, both isoforms interact with WDR68 (Fig. 1b). Interestingly, patients with disruptions of the C-terminal region of the AUTS2 locus tend to have more severe phenotypes than those with N-terminal disruptions (Beunders et al., 2013), indicating a PRC1-independent function carried out by the C-terminal domain of Auts2-S or/and Auts2-L.

**FIGURE 1.**
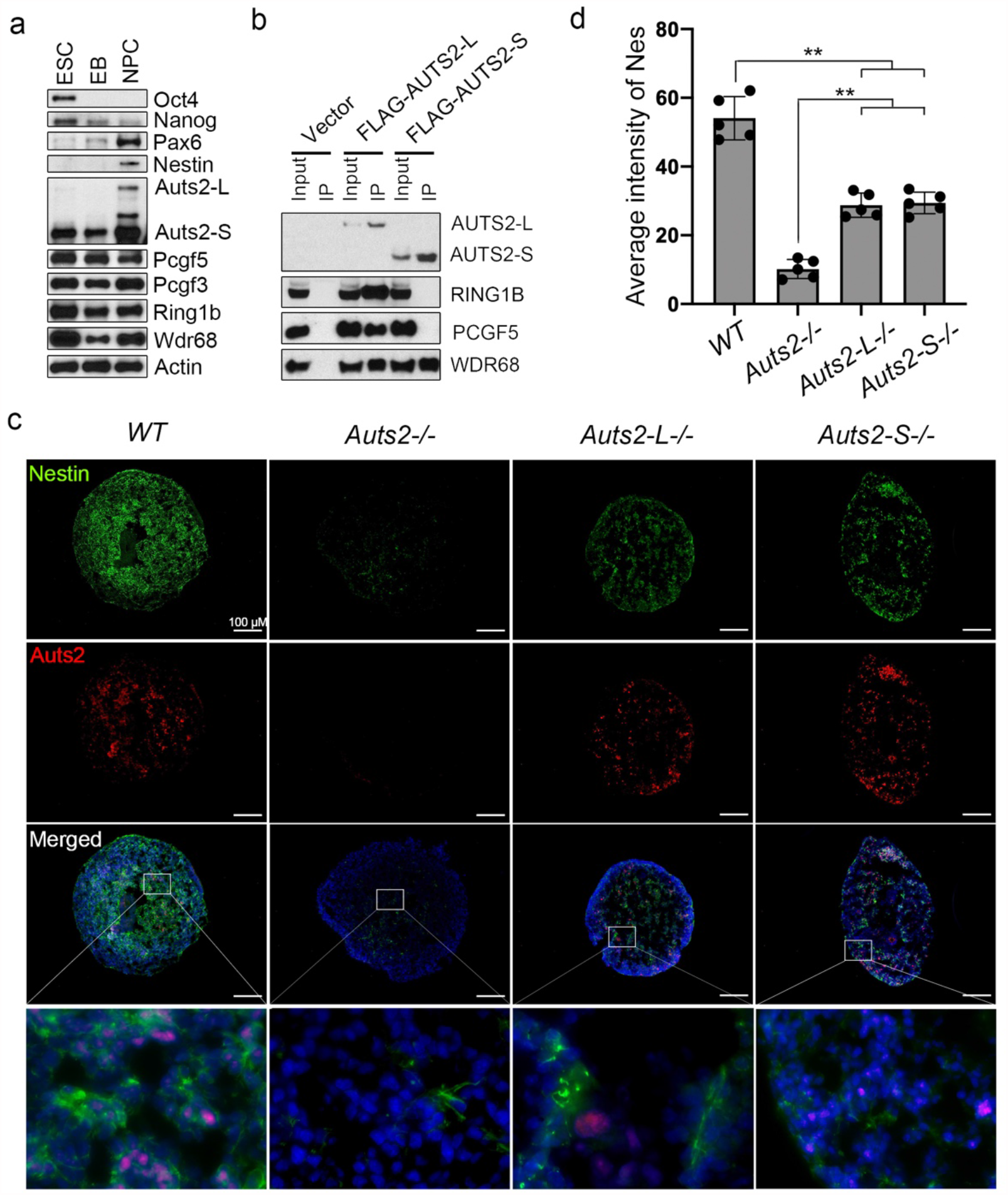
Auts2 is required for neuronal differentiation of mouse embryonic stem cells. ***a***, Immunoblotting of wild-type (WT) mouse embryonic stem cells (ESC), embryoid bodies (EB), and neuronal progenitor cells (NPC), using antibodies as indicated. ***b***, Immunoprecipitation (IP) with FLAG antibody-conjugated M2 beads from nuclear extract (NE) of HEK293T cells transfected with N-terminal FLAG and HA-tagged AUTS2 long-form (NFH-AUTS2-L), short-form (NFH-AUTS2-S), or control plasmid (Vector). Bound proteins were resolved on SDS-PAGE and detected by immunoblotting for the antigens indicated. 5% of input was loaded in all cases unless otherwise indicated. ***c***, Immunofluorescence staining of Nestin (Green) and Auts2 (Red) in mouse ESCs differentiated at NPC stage. DAPI is in Blue. The bottom panel shows the blowup of the box region as indicated. *Auts2-/-*, complete *Auts2* knockout; *Auts2-L-/-, Auts2-L* specific knockout; *Auts2-S-/-, Auts2-S* specific knockout. ***d***, Quantification of average Nestin immunofluorescence intensity by Image J. Each value is the mean of three independent measurements with error bars representing standard error. ** indicates *p*<0.01 by two-sided *t*-test.

To understand how these Auts2 isoforms affect neuronal differentiation, we used CRISPR/Cas9 mediated gene editing to engineer three different ESC lines, deleting Auts2-L, Auts2-S, or both (Fig. S1). Due to the shared coding region between Auts2-L and Auts2-S, it is impossible to use a simple CRISPR strategy that deletes or edits exons to create a null mutant specifically for Auts2-S without affecting Auts2-L. When analyzing our previous H3K4me3 ChIP-seq data, we found an H3K4me3 peak located just before exon 6 (Fig. S1a), indicating an internal promoter region for the alternative transcription start site. We reasoned that deletion of the intron region underlying this alternative promoter is likely to specifically silence Auts2-S (Fig. S1d). Indeed, using this strategy, we were able to generate an ESC line with only Auts2-S deleted (*Auts2-S-/-*), which was confirmed by genomic PCR, Sanger sequencing, and Immunoblotting (Fig. S1g, j, and k). In addition, we generated and validated ESCs with both isoforms (*Auts2-/-*) or only Auts2-L deleted (*Auts2-L-/-*) by targeting exons 9 and 7 respectively (*Auts2-/-*, Fig. S1b, e, h, and k; *Auts2-L-/-*, Fig. S1c, f, i and k).

To investigate the roles of Auts2-L and Auts2-S in neuronal cell fate determination, we performed immunofluorescence (IF) on the NPCs differentiated from various *Auts2* deficient ESCs. We found that the protein level of Nestin, an NPC marker, is almost completely diminished in *Auts2-/-*cells, whereas a significant reduction in *Nestin* expression was observed in *Auts2-L-/-* or *Auts2-S-/-*cells (Fig. 1c). It is unlikely that this defect results from a change in ESC status. Using an MTT cell proliferation assay, we confirmed a lack of any noticeable difference in cell growth among all three null ESC lines compared with wild-type (WT) ESCs (Fig. S2a). Furthermore, the pluripotency of mESCs lacking Auts2 was not changed, as measured by their alkaline phosphatase activity (Fig. S2b) and protein and mRNA levels of Oct4 and Nanog (Fig. S2c and S3a). The differentiation defect observed in Auts2 null cells was further confirmed by RT-qPCR analysis. In agreement with the previously mentioned Nestin IF analysis, NPC markers such as *Pax6, Nes, Neurod1*, and *Sox1*, are significantly reduced in Auts2-/-cells and are only partially decreased in either *Auts2-L-/-* or *Auts2-S-/-*cells (Fig. S3b). Given the fact that only Auts2-L interacts with PRC1 components, we suspect that the neuronal differentiation defect seen in these Auts2 deficient ESCs may involve a PRC1-independent pathway. Nonetheless, both isoforms of Auts2 contribute to the regulation of the differentiation of mESCs to NPCs.

### 2) Auts2 deletion causes imbalanced expression of genes required for the development of three germ layers

With next-generation sequencing (RNA-seq), we conducted a transcriptomic analysis of both WT and Auts2-/-in ESCs and NPCs. As shown in Fig. 2a, we have identified 4015 differentially expressed genes between WT and Auts2-/-NPCs. Among those, 1845 are down-regulated, and 2170 are up-regulated in Auts2-/-NPCs compared with WT. Among the down-regulated genes, many neuroectodermal marker genes were observed among the top differentially expressed genes, including *Nes, Sox1, Pax6, Pax2, Neurod1, Nog*, and *Tubb3* (Fig. 2a and b). A gene ontology (GO) analysis confirmed that these down-regulated genes in *Auts2*-/-NPCs are enriched in functional groups including “nervous system development”, “axon guidance”, “neuron migration”, “axonogenesis”, “neuron differentiation”, “glial cell differentiation”, “hippocampus development”, and “neuron fate commitment” (Fig. 2b, top panel). RNA-seq analysis also identified a group of genes that are not only up-regulated in *Auts2*-/-NPCs but, interestingly, are associated with mesoderm and endoderm development, including *Foxa1, Foxc1, Snai1, Tgfb2, Foxf1, Tgfb3, Fabp4, Gata4, Gata6*, and *Tnnt2* (Fig. 2a and b). Further GO analysis revealed functional aspects related to the regulation of mesoderm and endoderm lineage specification, including “angiogenesis”, “heart development”, “vasculogenesis”, “atrial cardiac muscle tissue morphogenesis”, and “liver development” (Fig. 2b, bottom panel).

**FIGURE 2.**
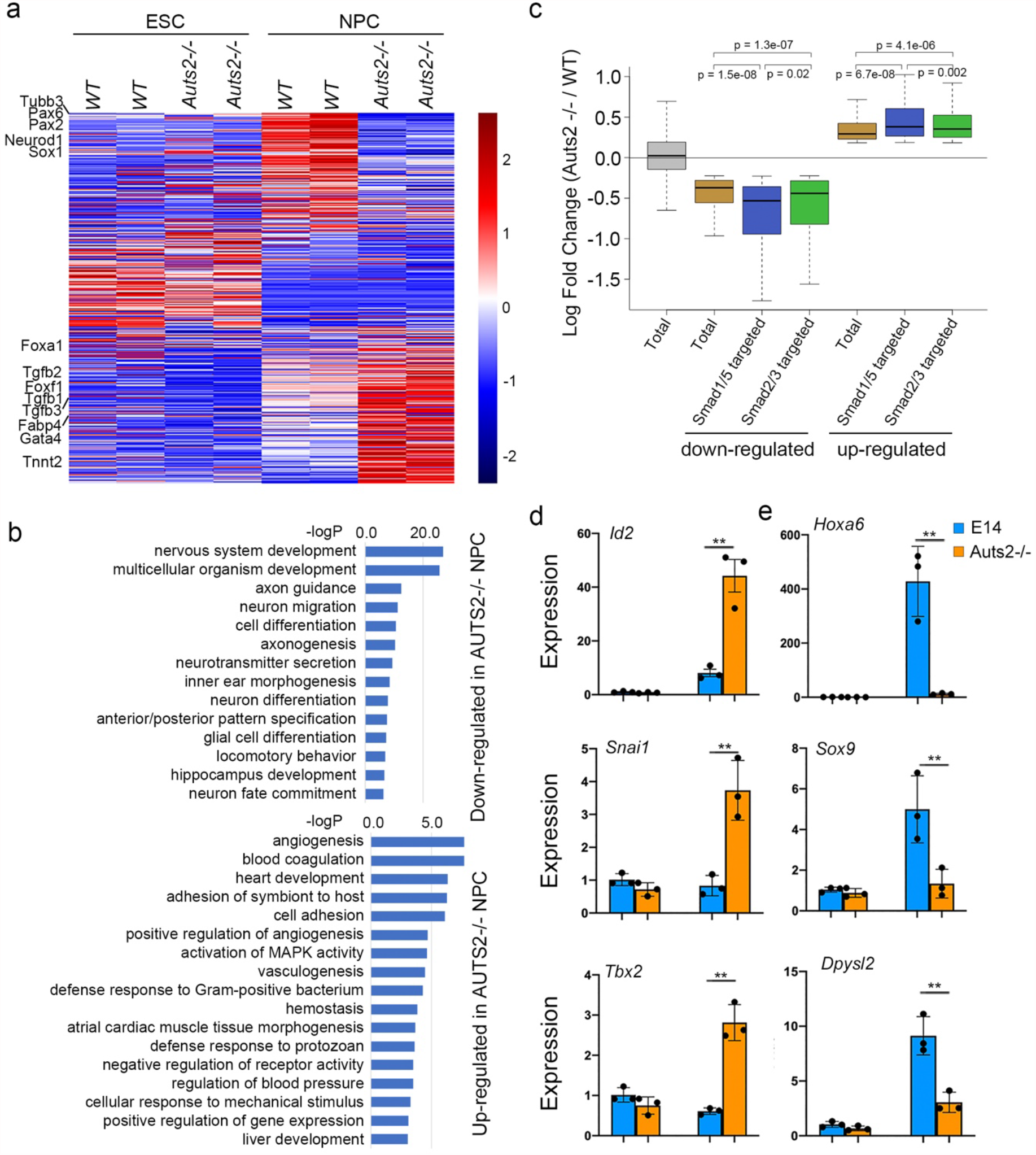
Auts2 controls the proper expression of lineage-specific genes. ***a***, Heatmap of transcriptomic analysis in WT and Auts2-/-ESCs and NPCs. RNA-seq analyses were performed on duplicate samples. The processed FPKM (reads per kilobase of exon per million reads mapped) values for each gene were used to calculate z scores used to generate the heatmap (See Supplemental Information for details). ***b***, GO analysis of genes that are down- or up-regulated in Auts2-/-NPCs compared with WT. The x-axis (in logarithmic scale) corresponds to the binomial raw P values. ***c***, Whisker-box plot shows the expression of genes targeted by SMAD1/5 or SMAD2/3. Fold change of RPKM values between WT and Auts2-/-NPCs from all expressed genes, down- or up-regulated genes, down- or up-regulated SMAD1/5-targeted genes, and down- or up-regulated SMAD2/3-targeted genes are plotted. SMAD1/5 and SMAD2/3 target genes are obtained from (Morikawa et al., 2011). ***d***, Expression of BMP responsive genes during differentiation, measured by quantitative RT-PCR. All mean values and standard deviations were calculated from three independent measurements. ** indicates *p*<0.01 by two-sided *t*-test.

The similarity in transcriptional signatures between cells with a dysregulated TGF-β/BMP pathway and *Auts2*-/-mESC leads to the assumption that Auts2 may regulate neuronal differentiation via the TGF-β /BMP pathway (Watabe and Miyazono, 2009). To better understand the influence of TGF-β/BMP signaling on the AUTS2-mediated regulation of neuronal differentiation, we took advantage of previously identified target genes for these pathways and examined their changes during differentiation in our WT and *Auts2*-/-NPCs. As shown in Fig. 2c, BMP-specific SMAD1/5 target genes that are also differentially regulated between WT and *Auts2*-/-NPCs tend to show more dramatic expression changes than all up-regulated or down-regulated genes. TGF-β-specific SMAD2/3 target genes exhibit a similar pattern but to a lesser extent (Fig 2c). With RT-qPCR, we further showed that BMP target genes are dysregulated in NPCs upon Auts2 deletion (Fig. 2d). These results strongly suggest that the BMP pathway is involved in Auts2-mediated regulation of neuronal differentiation in mouse ESCs.

### 3) Auts2 is required for inhibition of BMP signaling during neuronal differentiation

Upon stimulation, SMAD1/5/9, BMP-specific R-SMADs, are phosphorylated and dimerize with SMAD4, which is then translocated to the nucleus to regulate gene transcription. To understand how disruption of Auts2 may lead to the deregulation of the BMP pathway, we performed immunoblotting to examine the phosphorylation of SMADs. In keeping with our RNA-seq analysis, we found that the deletion of Auts2 results in an increase of phosphorylated Smad1/5/9 levels at the EB and NPC stages compared with WT cells, although no noticeable difference was seen in ESCs between *Auts2*-/- and WT (Fig. 3a). This observation further indicates that the effect of the loss of Auts2 on neuronal differentiation is due to up-regulated BMP signaling. The TGF-β pathway does not play a significant role as only a minimal increase of SMAD2 phosphorylation was found in *Auts2*-/-cells (Fig. 3a). To further examine the impact of Auts2 on acute cell response to BMP stimuli, we treated WT and *Auts2*-/-ESCs with BMP4 and evaluated the phosphorylation of Smad1/5/9. As expected and compared with WT cells, *Auts2*-/-cells displayed a significantly higher level of pSmad1/5/9 (Fig. 3b).

**FIGURE 3.**
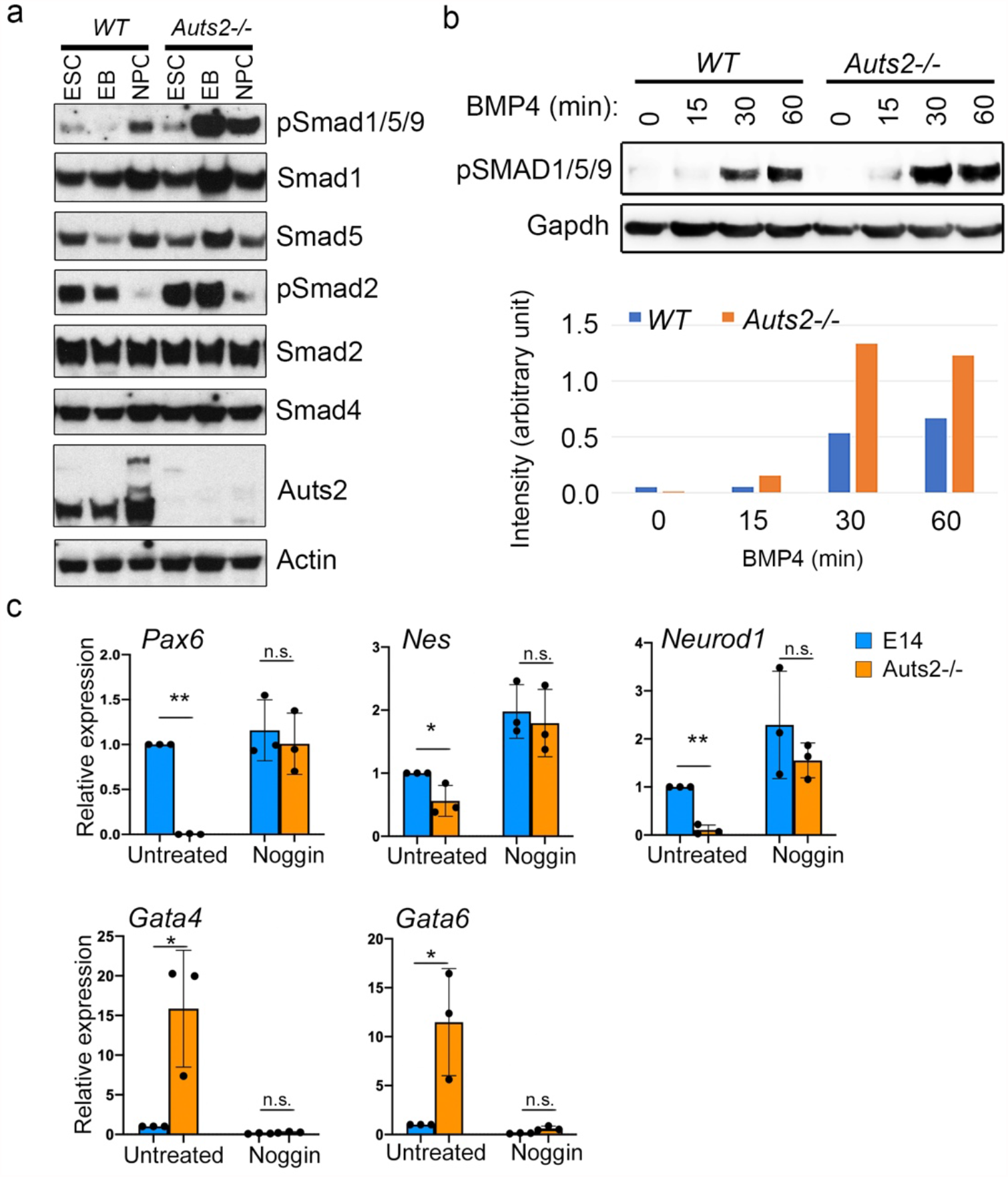
Auts2 is required for the inhibition of BMP signaling during neuronal differentiation. ***a***, Immunoblotting of WT and Auts2-/-mouse ESC, EB, and NPC using antibodies as indicated. ***b***, WT and Auts2-/-ESCs were stimulated with BMP4 at 25 μM for the time as indicated. Activation of BMP signaling was measured by the level of Smad1/5/9 phosphorylation by immunoblotting. ***c***, Expression of selected marker genes (neuroectoderm: Pax6, Nes, and Neurod1; mesoderm: Gata4; endoderm: Gata6) in Noggin-treated or untreated NPC of WT or Auts2-/-, measured by quantitative RT-PCR. All mean values and standard deviations were calculated from three independent measurements. * *p*<0.05, ** *p*<0.01, n.s. not significant.

The BMP pathway plays an essential role in cell differentiation. Studies have shown that activated BMP signaling promotes mesoderm and endoderm differentiation at the expense of neuroectoderm lineage (Li and Chen, 2013). To test whether up-regulated BMP signaling in *Auts2*-/-cells contributes to cell fate switch, we treated ESCs with noggin, an inhibitor of the BMP pathway, during differentiation. We found that noggin treatment rescued the dysregulation in marker genes of three germ layers (Fig. 3c). Viewing these results together, it is clear that Auts2 plays a critical role in repressing BMP signaling, serving as a crucial step for proper neuronal differentiation.

### 4) AUTS2 forms a neuronal lineage-specific complex with WDR68 and SKI

Both Auts2-L and Auts2-S interact with Wdr68 (Fig. 1b). Previously, we have shown that during NPC differentiation, the deletion of Wdr68 results in repression of the neuroectodermal markers, accompanied by the aberrant induction of mesodermal and endodermal markers (Wang et al., 2018). This observation is similar to our result in *Auts2*-/-cells (Fig. 1). Wdr68 belongs to a large family of WD40 domain-containing proteins that generally act as adaptors or scaffold factors for other proteins or protein complexes (Li and Roberts, 2001; Stirnimann et al., 2010). These observations suggest that Auts2-L or Auts2-S and Wdr68 may form a complex with unknown proteins that, in turn, may regulate BMP signaling during neuronal differentiation. To understand the molecular mechanism of Auts2-mediated control on neuronal cell fate, we performed proteomic analysis to identify novel binding partners for Auts2 and Wdr68. For this purpose, we used CRISPR/Cas9-mediated gene editing to generate an engineered ESC line in which tandem FLAG and HA tags were placed immediately in front of the translational start site of Wdr68 (E14: NFH-Wdr68, Fig. S4a). Genomic PCR and immunoblotting validated successful targeting (Fig S4b and c). We then performed quantitative mass spectrometry analysis on affinity-purified biological triplicate samples from E14: NFH-Wdr68 ESCs and their derived NPCs. As shown in the volcano blot in Fig. 4a, many known Wdr68-associated proteins (Gao et al., 2014; Miyata et al., 2014; Wang et al., 2018), including Pcgf3, Pcgf5, Rnf2, and Dyrk1a, were recovered, demonstrating the success of our proteomic analysis. In addition, Sloan–Kettering Institute (Ski), a previously identified inhibitor of the TGF-β/BMP pathway (Luo et al., 1999; Stroschein et al., 1999; Sun et al., 1999), was the most enriched in NPCs (Fig. 4a). Auts2 also was enriched in NPCs but less abundant compared with Ski (Fig. 4a). Immunoprecipitation in E14: NFH-Wdr68 cells further confirmed the association of Ski as well as Auts2-L and Auts2-S with Wdr68 (Fig. 4b). More interestingly, Ski was only immunoprecipitated from NPCs (Fig. 4b). This suggests that Wdr68-Ski interaction is highly specific to this stage. Meantime, both Auts2-L and Auts2-S are more highly expressed in NPC and more abundant in the immunoprecipitates (Fig. 4b). To examine if Auts2 actually interacts with Ski, we transfected HEK293T cells with long and short forms of N-terminal FLAG and HA-tagged AUTS2 (NFH-AUTS2-L and NFH-AUTS2-S) and performed immunoprecipitation with FLAG antibody-conjugated M2 beads. As expected, both forms of AUTS2 interacted with SKI and WDR68, but only AUTS2-L associated with PRC1 complex component PCGF5 (Fig. 4c). To further pinpoint the domains of AUTS2 required for interactions with different factors, we conducted immunoprecipitation experiments with various AUTS2 constructs. Based on these results, we found that the N-terminal fragment containing the first 208 amino acids is required for binding PRC1 components such as PCGF5 and RING1B, and the internal region between 600 to 700 amino acids was essential for binding WDR68 and SKI (Fig. 4d and Fig. S5). Through glycerol gradient analysis on FLAG affinity-purified complexes from a stable HEK293T-REx line expressing FLAG-HA-SKI, we observed that SKI, AUTS2, and WDR68 are present within the same fractions (Fig. 4e). This indicates the presence of a stable complex consisting of these three proteins. In summary, our proteomic and biochemical analyses defined a novel protein complex formed by Auts2, Wdr68, and Ski. Hereafter we will refer to this novel complex as the AWS complex (Fig. 4f). While both Auts2-L and Auts2-S are capable of forming the AWS complex, only Auts2-L can form the previously identified PRC1-AUTS2 complex (Fig. 4f) (Gao et al., 2014).

**FIGURE 4.**
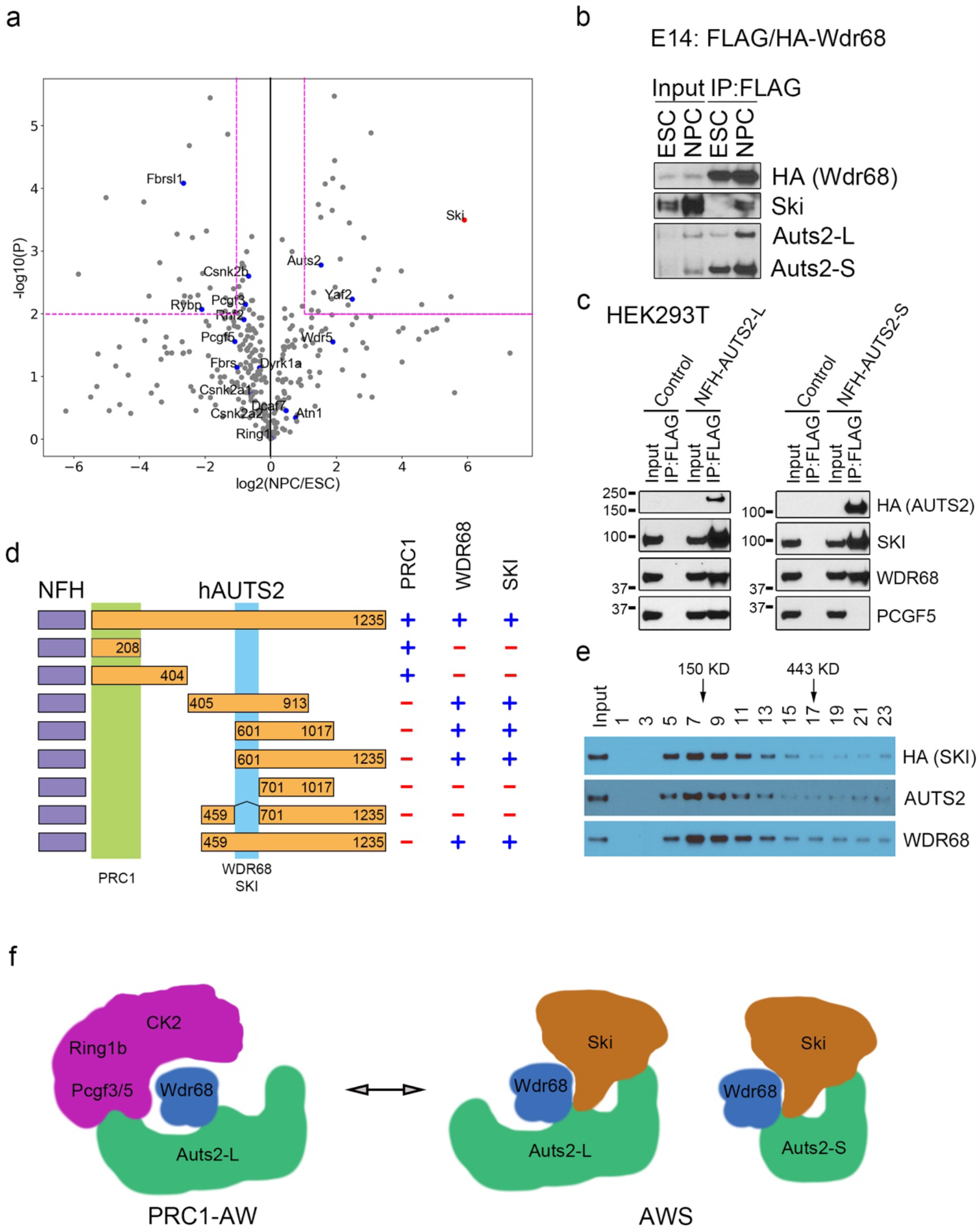
A novel protein complex formed by Auts2, Wdr68, and Ski. ***a***, Tandem affinity purification (TAP) followed by quantitative mass spectrometry analysis (MS) revealed a novel binding partner for Wdr68 in NPC, shown by a volcano blot. An E14 cell line with engineered Wdr68 locus inserted with N-terminal FLAG and HA tags (E14: NFH-Wdr68) was either cultured at ESC or differentiated to NPC stages. TAP was performed in triplicate samples of ESCs and NPCs, followed by label-free quantitative MS analysis (see Experimental Procedures). The X-axis shows the mean ratio of peptide intensity between NPC and ESC, and the Y-axis corresponds to P-value. ***b***, IP from NE of E14: NFH-Wdr68 cells at the ESC or NPC stages, using FLAG M2 beads. Bound proteins were resolved on SDS-PAGE and detected by Western blotting for the antigens indicated. ***c***, IP from NE of HEK293T cells transfected with NFH-AUTS2-L or NFH-AUTS2-S or vector control, using FLAG M2 beads. Bound proteins were resolved on SDS-PAGE and detected by Western blotting for the antigens indicated. ***d***, Mapping of AUTS2 domains required for interaction with indicated proteins or protein complexes. Plasmids expressing NFH-AUTS2 with various lengths were expressed in HEK293T cells followed by IP with M2 beads to detect their interaction with PRC1 components, WDR68 and SKI, as indicated on the right (See Experimental Procedures for details). The deduced domains required for specific interaction are highlighted and indicated at the bottom. ***e***, Glycerol gradient (15-35%) analysis of FLAG-purified NFH-AUTS2-S (See Experimental Procedures). Every other fraction was resolved on SDS-PAGE followed by immunoblotting for the antigens indicated. The fractions containing AUTS2, WDR68, and SKI simultaneously indicate the presence of a complex formed with these proteins (AWS). ***f***, Schematic model of AUTS2-containing complexes. See text for details.

### 5) AWS negatively regulates the stability of phosphorylated SMAD1/5/9 through CUL4-mediated poly-ubiquitination

SKI has been previously identified as an inhibitor of TGF-β/BMP signaling by directly binding to the R-SMAD/co-SMAD complex (Luo et al., 1999), thus unlikely to affect the level of R-SMAD phosphorylation level. Furthermore, transcriptional feedback does not likely explain this phenomenon as the enhancement of pSmad in *Auts2*-/-ESCs was seen shortly after 15 min of BMP stimulation (Fig. 3b). Another possibility is that the AWS complex may affect the activation step of SMAD signaling. However, we ruled out this explanation since Auts2 is predominantly expressed in the nucleus (Bedogni et al., 2010; Gao et al., 2014).

We then asked whether the AWS complex affects the stability of SMAD1/5/9. It has been previously shown that WDR68 is present in the CUL4 E3 complex as a substrate receptor that directs specific substrate binding for polyubiquitination and subsequent proteasomal degradation (Higa et al., 2006; Jin et al., 2006). WDR68 directly interacts with Damage Specific DNA Binding Protein 1 (DDB1), an adaptor specific to the CUL4 E3 complex (Jin et al., 2006). Using immunoprecipitation, we demonstrated that AUTS2 also interacts with DDB1 (Fig. 5a). Therefore, we hypothesized that the AWS complex might affect the stability of the pSMAD by mediating its polyubiquitination. Indeed, when we transfected HEK293T cells with HA-ubiquitin (HA-ub) and FLAG-SMAD followed by immunoprecipitation with an HA antibody, we recovered polyubiquitinated SMAD1, which was further enhanced by treatment with MG132, a proteasome inhibitor (Fig. 5b). More interestingly, when these cells were treated with a pan CUL E3 inhibitor, MLN4924, polyubiquitinated SMAD1 was reduced (Fig. 5b), indicating that CUL E3 contributes to the polyubiquitination of SMAD1. Consistent with this observation, the addition of MLN4924 results in increased pSMAD1/5/9 upon BMP4 treatment in HEK293T cells (Fig. 5c). To further examine if the effect on pSMAD1/5/9 is due to CUL4 specifically, we silenced the CUL4-specific adaptor DDB1 using siRNA and found that the DDB1 knockdown led to elevated SMAD activation compared with the control knockdown (Fig. 5d). Similarly, exogenous expression of a dominant-negative form of CUL4 (DN-CUL4) led to up-regulation of pSMAD1/5/9 (Fig. 5e). To further test our hypothesis, we transfected HEK293T cells with plasmids to express AWS components and examined their effects on the pSMAD induced by BMP treatment. As shown in Fig. 5f, overexpression of AUTS2 or SKI led to a reduction in pSMAD in response to BMP4 compared with the control transfection, whereas co-transfection of AUTS2 and SKI caused a slightly more substantial decrease of pSMAD. Interestingly, total SMAD levels remained unchanged (Fig. 5f), suggesting that the effect of AWS was specific to pSMAD. Taken together, our biochemical analyses delineated a regulatory pathway for restricting BMP-specific SMAD signaling by the AWS and CUL4 E3 complex, therefore promoting proper neuronal differentiation.

**FIGURE 5.**
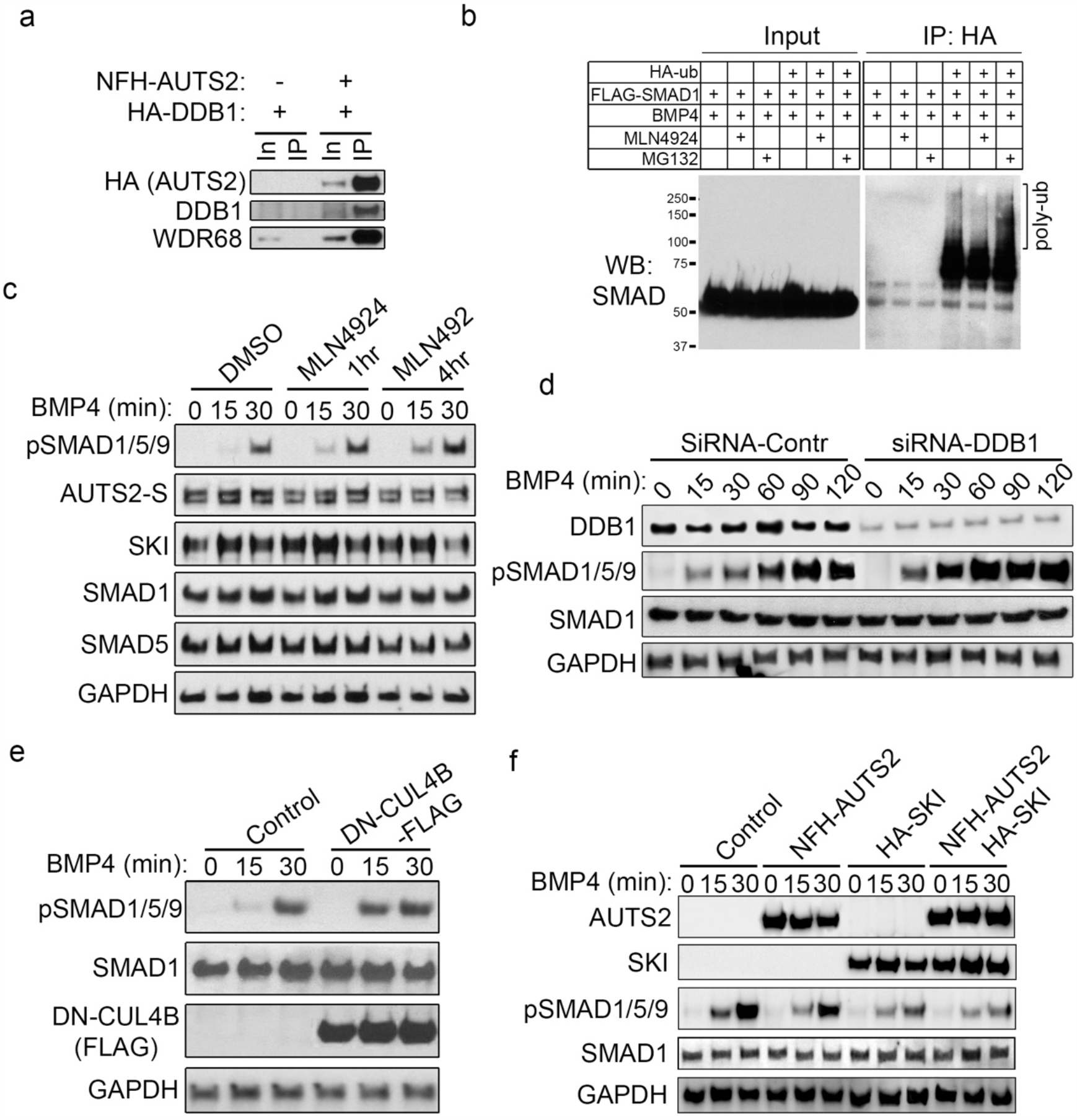
The AWS complex promotes degradation of pSMAD1/5/9 through CUL4 E3. ***a***, IP in HEK293T cells transfected with NFH-AUTS2 and HA-DDB1, using FLAG M2 beads. Bound proteins were resolved on SDS-PAGE and detected by Western blotting for the antigens indicated. ***b***, *in vivo* poly-ubiquitination assay. HEK293T cells were transfected with HA-ub and FLAG-SMAD1. Two days after transfection, cells were lysed in the presence of 1% SDS and immunoprecipitated with HA beads, followed by immunoblotting with SMAD1 antibody. ***c***, Cullin E3 negatively regulates the level of pSMAD1/5/9. HEK293T cells were stimulated with BMP4 at various times as indicated with or without prior treatment with MLN4924, a pan inhibitor for CUL E3, followed by immunoblotting with antibodies as indicated. ***d***, Two days after treatment of control or siRNA for DDB1, HEK293T cells were stimulated with BMP4 at various times as indicated, followed by immunoblotting. ***e***, Two days after transfection with a dominant-negative form of CUL4 (DN-CUL4B), HEK293T cells were stimulated with BMP4 at various times as indicated, followed by immunoblotting. ***f***, HEK293T cells were transfected with AUTS2 and SKI, individually or combinatorically, then stimulated with BMP4, followed by immunoblotting.

### 6) Auts2 control of neuronal differentiation requires the formation of the AWS complex

To understand how the interaction of Auts2 with Wdr68 and Ski affects its function during neuronal differentiation, we generated a lentivirus expressing either hAUTS2-S or a version with deleted amino acids 600-700 (hAUTS2-S-Δ). According to our domain mapping results (Fig. 4d), hAUTS2-S-Δ does not interact with Wdr68 and Ski and is therefore incapable of forming the AWS complex. We infected WT and *Auts2*-/-ESCs with these same lentiviruses; these ESCs were then differentiated to NPCs and scrutinized using RNA-seq analysis. We first examined whether the infection was successful by detecting RNA-seq tags of hAUTS2-S and hAUTS2-S-Δ We found that RNA-seq signals covered all exons from 8 to 19 in hAUTS2-S infected cells, but in hAUTS2-S-Δ infected cells, there was no coverage for exons 11-14, which corresponds to translated amino acids 600-700 that were deleted (Fig. 6a). Next, we analyzed the global transcriptomic change between WT NPCs with mock infection (WT, mock) and *Auts2*-/-NPCs with infection of either mock, hAUTS2-S, or hAUTS2-S-Δ (*Auts2*-/-, mock/ hAUTS2-S/ hAUTS2-S-Δ). While the reintroduction of hAUTS2-S in *Auts2*-/-NPCs blocks the transcriptional deregulation in both directions compared with mock-infected WT NPCs (Fig. 6b, top and middle panels), expression of hAUTS2-S-Δ in *Auts2*-/-NPCs failed to do the same (Fig. 6b, top and bottom panels). Furthermore, down-regulation of neuroectodermal marker genes, including *Sox1, Neurod1, Pax6, Map2, and Tubb3* and up-regulation of mesodermal and endodermal marker genes, including *Hand1, Twist2, Snai1, Gata4*, and *Gata6*, are both rescued in hAUTS2-S infected NPCs but not in hAUTS2-S-Δ infected NPCs (Fig. 6b). Based on our clustering analysis, the overall gene expression signature of hAUTS2-S infected NPCs is closer to mock-infected WT NPCs, while hAUTS2-S-Δinfected NPCs were more similar to mock-infected *Auts2*-/-NPCs (Fig. 6c).

**FIGURE 6.**
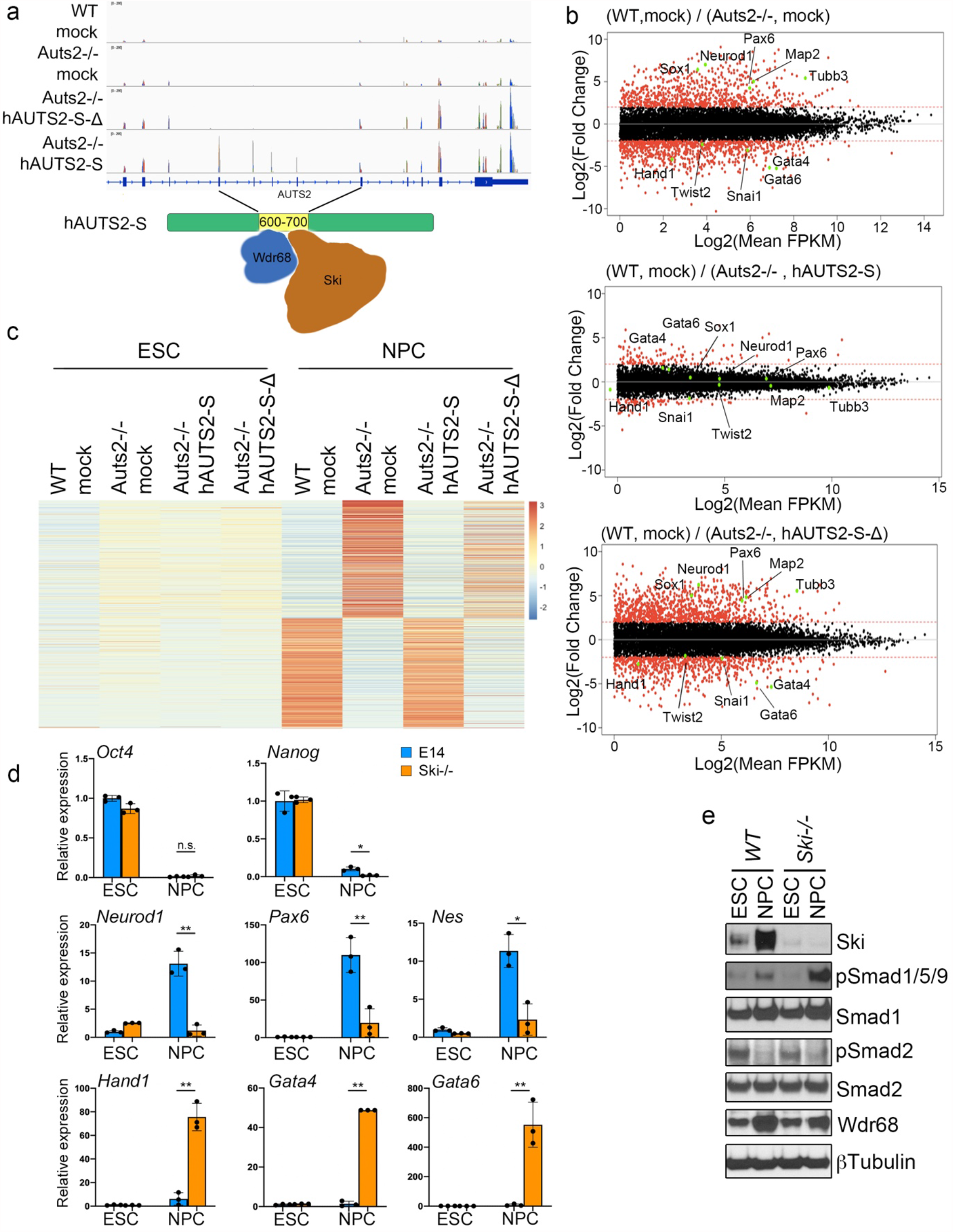
Ski mediates Auts2 regulation on neuronal differentiation. ***a***, RNA-seq reads confirm the expression of human-specific short form AUTS2 (hAUTS2) with or without deletion between 600 to 700 amino acids (hAUTS2-S-Δ and hAUTS2-S) through lentiviral infection in *Auts2*-/-mouse ESCs. A schematic of the deleted region of AUTS2 that promotes AUTS2 - WDR68 and AUTS2 -SKI interaction is shown at the bottom. ***b***, Comparison of gene expression levels based on RNA-seq analysis from WT and *Auts2*-/-NPCs with lentiviral infection with mock, hAUTS2-S and hAUTS2-S-¹’. The x-axis is the log_2_ value of the average FPKM of a gene and the y-axis is the log2 value of fold changes of FPKM of a gene between two groups. Genes with an average FPKM below one are filtered out. ***c***, Heatmap of transcriptomic analysis in WT and *Auts2*-/-ESCs and NPCs, with lentiviral infection as indicated. Z scores were calculated using processed FPKM values for each gene as in Fig. 2a. ***d***, Expression of selected marker genes (pluripotency: *Oct4* and *Nanog*; neuroectoderm: *Pax6, Nes*, and *Neurod1*; mesoderm: *Hand1* and *Gata4*; endoderm: *Gata6*) in ESC or NPC of WT or *Ski*-/-, measured by quantitative RT-PCR. All mean values and standard deviations were calculated from three independent measurements. * *p*<0.05, ** *p*<0.01, n.s. not significant. ***e***, Immunoblotting of WT and *Ski*-/-mouse ESC and NPC using antibodies as indicated.

Ski has been previously shown to regulate brain development in mice (Berk et al., 1997), but the underlying mechanism is not fully understood. Our results suggest a novel role for Ski in regulating neuronal differentiation by promoting degradation of pSMAD1/5/9, which depends on its interaction with Auts2 and Wdr68. We generated ESC lines with deleted *Ski* (*Ski*-/-) through CRISPR to further test this hypothesis. Similar to *Auts2*-/-ESCs, *Ski*-/-ESCs showed defects in differentiation to NPCs, as evidenced by RT-qPCR analysis (Fig. 6d). More interestingly, this differentiation defect is accompanied by the up-regulation of pSMAD1/5/9 signaling (Fig. 6e). In summary, our results demonstrate that Auts2 promotes neuronal differentiation through working together with Wdr68 and Ski in a protein complex.

### 7) The AWS complex is required for proper gene expression and BMP signaling in mouse cortical neurons

To investigate the role of the AWS complex *in vivo*, we turned to a mouse model with CNS-specific deletion of both Auts2 isoforms through targeting exon 9 (Fig. 7a, gift from Danny Reinberg). As previously described, Auts2 is highly enriched in the neocortex of mice embryos and the cortical neurons in adult mice. Therefore, we isolated and cultured cortical neurons from WT and Auts2+/-mice at E15. Unfortunately, we were not able to obtain *Auts2*-/-mice due to potential embryonic lethality. Through RT-qPCR analysis, we found that, compared with WT, *Auts2*+/-primary cortical neurons showed lower expression of neuronal markers (Fig.7b). In agreement with our *in vitro* analysis on mouse ESC and NPC (Fig. 2a and b), mesodermal and endodermal markers were also aberrantly induced in *Auts2*+/-neurons (Fig. 7c).

**Figure 7.**
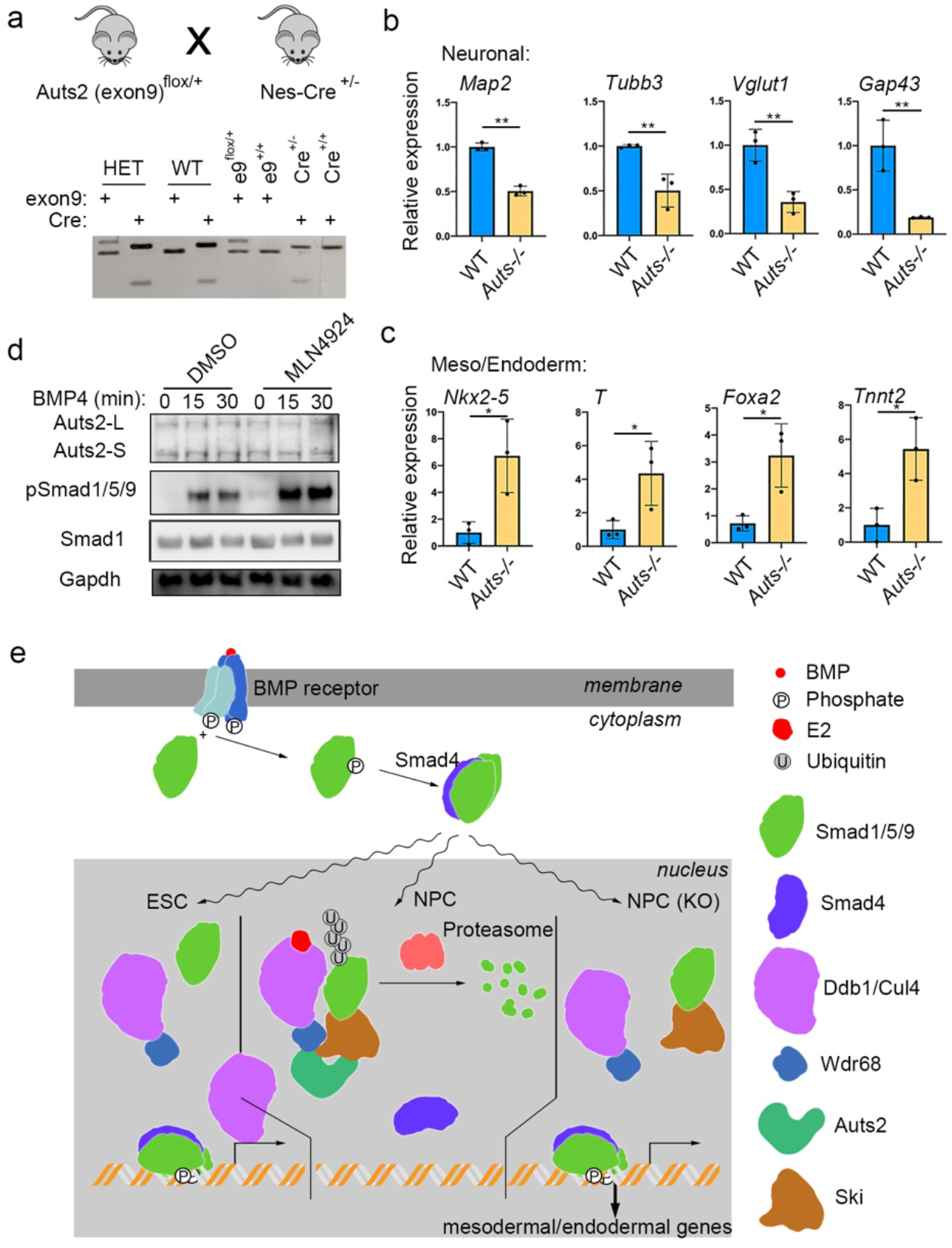
Auts2 is required for proper gene expression and BMP signaling in mouse primary cortical neurons. ***a***, Genotyping PCR in WT and *Auts2*+/-mouse primary cortical neurons. ***b***, Relative expression of neuron markers between WT and *Auts2*+/-mouse cortical neurons. ***c***, Relative expression of mesoderm and endoderm marker genes. (***b*** and ***c***) Each value is the mean of three independent measurements with error bars representing standard error. * *p*<0.05, ** *p*<0.01. ***d***, WT cortical neurons were treated with MLN4924 or DMSO for 1 hour, then treated with BMP4 for 0, 15, and 30 minutes, followed by immunoblotting. ***e***, Model of AWS/CUL4 E3-mediated regulation of BMP signaling during neuronal differentiation. See text for details.

Mechanistically, our ESC studies discovered the AWS-mediated negative regulation on BMP signaling through CUL4-mediated polyubiquitination and proteasomal degradation of pSMAD1/5/9 during NPC differentiation. To test if this mechanism plays a role in brain development, we treated the WT cortical neurons with either DMSO or MLN4924, followed by the acute stimulation by BMP4. As shown in Fig. 7d, pSMAD1/5/9 was increased upon the stimulation of BMP signaling with CUL inhibition (MLN4924), compared with control (DMSO). In summary, it is highly likely the AWS complex plays a role in restricting BMP signaling during neuronal differentiation *in vivo*. Furthermore, combining our mouse ESC and mouse cortical neuron studies, we uncovered a novel pathway through which AUTS2, an ASD/NDD risk factor, forms an NPC-specific complex and employs the CUL4 E3 complex to regulate BMP signaling, thereby affecting the cell fate determination in neurodevelopment (Fig. 8).

## Discussion

Our mechanistic dissection of AUTS2-mediated regulation of neuronal differentiation led to discovering a novel protein complex formed by AUTS2, WDR68, and SKI (referred to as AWS), impacting BMP signaling. To maintain proper cell fate choice during development, the BMP pathway needs to be tightly regulated; its inhibition is vital for engaging a transcriptional program that promotes neuronal differentiation (Li and Chen, 2013). Through biochemical analyses, we demonstrate that the AWS complex, which is formed specifically in the neural progenitor stage, targets BMP-specific SMADs to the CUL4 E3 complex for poly-ubiquitination and proteasomal degradation. Indeed, loss of Auts2 results in an up-regulation of BMP signaling and severe defects in neuronal differentiation in mouse ESCs, which is rescued by inhibition of the BMP pathway. These results uncover a novel mechanism through which a prominent NDD risk factor controls neural differentiation, providing a new opportunity for therapeutic interventions.

Although disruption of *AUTS2* was initially identified in autism patients, subsequent genetic and genomic studies revealed that *AUTS2* deregulation is linked to various forms of NDD (Oksenberg and Ahituv, 2013). The phenotypic variability created by *AUTS2* disruptions is accompanied by distinct transcriptional isoforms, but the underlying molecular mechanisms are not understood. It has been observed that patients with exonic deletions in the 3’
s region of the *AUTS2* locus tend to have more severe phenotypes than those having 5’ disruptions (Beunders et al., 2013). An alternative transcription start site prior to exon 6 drives the expression of a short isoform of *AUTS2* (*AUTS2-S*), which is in frame with the long isoform (*AUTS2-L*). In the present study, we found that both AUTS2-L and AUTS2-S, via their shared middle region, form the AWS complex by associating with WDR68 and SKI, whereas AUTS2-L forms a specific PRC1-AUTS2 complex depending on its N-terminus (Fig. 4). Targeting exon 9 of Auts2 in mouse ESCs, which results in the loss of both isoforms, clearly shows more dramatic defects in neuronal differentiation than the specific deletion of Auts2-L. These results echo the severity of dysmorphic phenotypes in patients with disruptions of AUTS2 at different regions (Oksenberg and Ahituv, 2013) and indicate the potential involvement of a PRC1-independent pathway affected by these genetic altercations. Aberrations of BMP signaling have been closely linked to several craniofacial dysmorphic syndromes (Graf et al., 2016), which is highly likely the mechanism shared by both AUTS2 isoforms. However, given the many other factors associated with AUTS2 (Gao et al., 2014; Monderer-Rothkoff et al., 2021), further research is needed to elucidate the interplay between PRC1-dependent mechanism, BMP signaling, and maybe other pathways to understand how exactly AUTS2 regulates neurodevelopment.

Previously, the targeted deletion of *Ski* in mice led to developmental defects in the CNS, most likely by regulating the fate of neuroepithelial stem cells (Berk et al., 1997). The TGF-β/BMP pathway consists of an intricate network of factors for both stimulatory and inhibitory regulation of signal transduction (Massague, 2012). SKI has been identified as a negative regulator of TGF-β/BMP signaling, and subsequent studies suggest that SKI interacts with SMAD complexes to keep them inactive, thereby preventing transcriptional engagement of downstream target genes (Luo et al., 1999). Our studies suggest a novel mechanism by which SKI (part of the AWS complex and via recruitment of CUL4 E3) engages the BMP pathway. This occurs during neuronal differentiation and subsequently leads to the removal of pSMAD1/5/9 through ubiquitination and proteasomal degradation, a process that promotes a transcriptional program that favors a specific neuronal trajectory. It will be interesting to see how the two modes of SKI inhibition on TGF-β/BMP signaling, the SKI-SMAD inert complex and the AWS/CUL4-mediated degradation of SMADs, coordinate to regulate neuronal differentiation in a developmental stage- or cell type-specific manner. A possible molecular switch may reside in a molecule like AUTS2 or other NDD risk factors based on our results. Future explorations into these avenues will likely provide mechanistic insight into the factors responsible for the fine-tuned regulation of these critical signaling pathways and their influence on cell fate choice during neurodevelopment. In addition, given the broad range of impact by BMP signaling on the development of various tissues, our discoveries may inspire a more specific targeting approach to correct the pathologically deregulated BMP pathway in NDD patients.

## Supporting information

Supplemental Infomation

## Author contributions

Z.Geng, Q.W., W.M., T.W., J.C., and E.G. conducted the experiments; R.H. and D.D. provided guidance and help on the immunofluorescence analysis; Z. Gao, J.S., and Y.W. designed the experiments; Z.Geng and Z.Gao wrote the paper. All authors contributed to the discussion of the manuscript.

## Acknowledgments

We would like to thank Dr. Kathleen Mulder for the discussion of the experiments. We thank Dr. Abraham Thomas and the Microscopy Imaging Core for assistance in imaging and Dr. Yuka Imamura and the Genome Sciences Facility for assistance in deep sequencing. The *Auts2* mouse strain is a gift from Dr. Danny Reinberg. This work was supported by the following NIH grants: R35GM133496 to Z. Gao; R00AA024837 to J. Stafford; R35ES031707 to Y. Wang.

## Experimental Procedures

### Cell culture

HEK293 T-REx cells expressing FLAG-HA-AUTS2-S were generated by transfection with pINTO-NFH-AUTS2-S (Table S1), followed by Zeocin selection. These cells and normal HEK293T cells were maintained in a standard DMEM medium containing 10% FBS (Atlanta Biologicals, Cat# S11050), L-glutamine, and penicillin/streptomycin. E14 mouse ES cells or their derived CRISPR-engineered cells were cultured in DMEM medium, supplemented with 15% FBS (ES certified, Atlanta Biologicals, Cat# S10250), LIF, non-essential amino acids, β-mercaptoethanol, L-glutamine, penicillin/streptomycin, sodium pyruvate, and two small-molecule kinase (MEK and GSK3) inhibitors (PD0325901, Cayman Chemical, Cat# 13034 and CHIR99021, Cayman Chemical, Cat# 13122). E14 cells were either cultured on MEF feeder cells or gelatin-coated plates.

### CRISPR/Cas9-mediated gene editing

Oligos corresponding to candidate sgRNA sequences were purchased from IDT and cloned into pX458 (Addgene) as previously described (Cong et al., 2013). Briefly, E14 cells were transfected with plasmids carrying sgRNAs with Lipofectamine (Life Technologies). After two days, transfected cells were sorted into 96-well plates coated with gelatin at one cell per well. Clones were tested by genomic PCR. Positive clones were further validated by Sanger sequencing and immunoblotting.

The sgRNA sequences used were (PAM trinucleotides are underlined):

*Auts2*-/-: mAuts2-KO-1: AGGTGCTGGCGTCGGCATGATGG mAuts2-KO-2: AATGCTGGGGCGCATCCCGTAGG

*Auts2-L*-/-: mAuts2-L-KO-1: GCGTATATTCCCTAAACTATGGG mAuts2-L-KO-2: CCACAGGGTAGGGTTACCATTGG

*Auts2-S*-/-: mAuts2-S-KO-1: GTCTGACATGGATGGGAGGTTGG mAuts2-S-KO-2: GAATGGTCACTAGCAAGCCTCGG

*Ski*-/-: mSki-KO-2: GGGCGGCCCGGCCGCTTTCTCGG mSki-KO-3: AGTCGCGCAGCACCGAGTTGAGG

NFH-mWdr68: mWdr68-KI: TCTACAAATATGAAGCGCCCTGG

### mESC neuronal differentiation

mESC differentiation was performed as previously described (Bibel et al., 2004). mESCs cultured on MEF feeders were transferred onto gelatin-coated plates for two passages before differentiation. Embryonic bodies (EB) were formed by culturing cells in differentiation medium (DMEM medium, 15% FBS, non-essential amino acids, β-mercaptoethanol, L-glutamine, penicillin/streptomycin, sodium pyruvate) in ultra-low attachment plates. After four days, retinoic acid was added to the medium at 5 μM for four more days to generate neuronal progenitor cells (NPC).

### Immunofluorescence

Differentiated EBs were fixed with 4% paraformaldehyde at room temperature for 20 min, followed by triple washing in PBS. Tissues were stocked in 30% PBS-buffered sucrose solutions overnight at 4 °C. EBs were embedded in O.C.T compound (Sakura) and cryosectioned at 20 μM thickness. For immunofluorescence imaging, sections were blocked and permeabilized in 0.5% Triton X-100 and 5% BSA in PBS. Sections were then incubated with an anti-Nestin primary antibody in 0.1% Triton X-100 and 1% BSA in PBS for 2 hours at room temperature. After washing 3 times in PBS, sections were incubated with Alexa Fluor 488 anti-mouse antibody (Gibco) for 1 hour at room temperature. Slides were washed 3 times, stained with DAPI, and mounted for imaging.

For image quantification, the fluorescence images were loaded and converted to greyscale in Image J. EB sections were selected by using drawing tools. The integrated fluorescence intensity of sections was measured by Image J. For each image, three random regions outside EB sections were measured and averaged, and used as background fluorescence intensity. The corrected fluorescence intensity of each EB was equal to the integrated intensity subtracted by background intensity.

### RNA-seq and analysis

RNA-seq was performed and analyzed as previously described (Gao et al., 2014). Briefly, cDNA libraries were prepared using the NEXTflex™ Illumina Rapid Directional RNA-Seq Library Prep Kit (BioO Scientific) as per the manufacturer’s instructions. Libraries were loaded onto a TruSeq Rapid flow cell on an Illumina HiSeq 2500 (located at the Genomic Sciences Facility at the Penn State University College of Medicine). These cells were run for 50 cycles using either a single-read or pair-end recipe according to the manufacturer’s instructions. Raw reads were aligned to the Mus musculus genome (UCSC mm10) using TopHat (version 2.1.1) (Trapnell et al., 2009) with default parameters. Subsequently, the relative abundances (FPKM) of the transcripts and genes were estimated with Cufflinks (version 2.2.1) (Trapnell et al., 2010). Low-expressed genes were filtered out and differentially expressed genes were identified by comparing the z-score difference at the NPC stage. Gene ontology (GO) analysis was performed using DAVID (Huang da et al., 2009). The z-scores of gene expressed matrix were used for heatmap plotting.

### Quantitative RT-PCR

Total RNA was extracted with TriPure reagent (Roche, Cat# 11667165001) and used to synthesize cDNA with the SuperScript III system (Invitrogen, Cat# 18080-044). The resulting cDNA was mixed with Brilliant III Ultra-Fast SYBR QPCR master mix (Agilent Cat#600883) and primers, then run on a Biorad CFX Connect real-time PCR detection system.

### Affinity purification

Affinity purification in ESCs or NPCs was performed in triplicates previously described (Gao et al., 2012). Approximately 10^8^ Cells were washed with PBS and then suspended in 10 ml Buffer A (10 mM Tris-HCl, pH 7.9, 1.5 mM MgCl2, 10 mM KCl, 0.5 mM DTT, 0.2 mM PMSF, 1 μg/ml Pepstatin A, 1 μg/ml Leupeptin, 1 μg/ml Aprotinin), incubated on ice for 30 min, and then homogenized by a 40 ml dounce homogenizer for 10 strokes, using the loose pestle. The suspension was then subjected to 15, 000 × g centrifugation at 4 °C for 10 min. The resulting nuclear pellet was resuspended in 10 ml Buffer C (20 mM Tris-HCl, pH 7.9, 25% glycerol, 420 mM NaCl, 1.5 mM MgCl2, 0.2 mM EDTA, 0.5 mM DTT, 0.2 mM PMSF, 1 μg/ml Pepstatin A, 1 μg/ml Leupeptin, 1 μg/ml Aprotinin), and subjected to another 10 strokes by the dounce homogenizer. After rotation at 4 °C for 30 min, the suspension was centrifuged at 40, 000 × g at 4 °C for 30 min, and the supernatant was nuclear extract (NE). 8 ml NE was mixed with 2 ml Buffer A, 0.02% NP-40, and 200 μl pre-washed FLAG M2 beads. After rotation at 4 °C overnight, the M2 beads were spun down at 2000 × g at 4 °C for 10 min. The beads were washed with Buffer W (2/3 volume of Buffer C, 1/3 volume of Buffer A, NP-40 0.02%) 5 times and then eluted with 500 μl FLAG peptides of 250 μg/ml in Buffer W by rotating at 4 °C for 1 hour. The M2 eluate was then incubated with 30 μl HA beads at 4 °C for 4 hours. The HA beads were washed with Buffer W 5 times and eluted by 100 μl glycine (0.1 M, pH 2.0), and then neutralized by adding 6.5 μl Tris solution (1.5 M, pH 8.8), resulting in the final HA eluate.

### Quantitative proteomic analysis

For quantitative proteomic analysis, the TAP enriched proteins were digested with modified MS-grade trypsin (Thermo Pierce) at an enzyme/substrate ratio of 1:100 in 50 mM NH4HCO3 (pH 8.5) at 37 °C for overnight. The resulting peptide samples were then loaded at 3 µL/min onto a precolumn (150 µm i.d.) comprised of a 3.5-cm column packed with 5 µm C18 120 Å reversed-phase material (ReproSil-Pur 120 C18-AQ, Dr. Maisch), for LC-MS/MS analysis. The trapping column was connected to a 20-cm fused silica analytical column (PicoTip Emitter, New Objective, 75 µm i.d.) with 3 µm C18 beads (ReproSil-Pur 120 C18-AQ, Dr. Maisch). The peptides were then separated using a 180-min linear gradient of 2-45% acetonitrile in 0.1% formic acid and at a flow rate of 250 nL/min. The mass spectrometer was operated in a data-dependent scan mode. Full-scan mass spectra were acquired in the range of m/z 350-1500 using the Orbitrap analyzer with a resolution of 70,000. Up to 25 most abundant ions found in MS with a charge state of 2 or above were sequentially isolated and collisionally activated in the HCD cell with a collision energy of 27 to yield MS/MS.

Maxquant (Cox and Mann, 2008), Version 1.5.2.8, was used to analyze the LC-MS and MS/MS data for the identification and quantification of proteins in the LFQ mode. The maximum number of miss-cleavages for trypsin was two per peptide. Cysteine carbamidomethylation was set as a fixed modification. Methionine oxidation and phosphorylation on serine, threonine, and tyrosine were set as variable modifications. The tolerances in mass accuracy for MS and MS/MS were both 20 ppm. Maximum false discovery rates (FDRs) were set to 0.01 at both peptide and protein levels, and the minimum required peptide length was six amino acids.

### Isolation and culture of mouse cortical neurons from Auts2 cKO mice

Generation of the conditional knockout of all Auts2 isoforms in the brain was performed by crossing mice harboring loxP sites flanking exon 9 of Auts2 with Nestin-Cre driver mice (Jackson Labs). Littermates were selected for the wild-type and heterozygous Auts2 knockout pups to match all conditions for subsequent experiments. Primary cortical neurons were prepared from postnatal day 0 mouse pups by first dissecting the individual mouse pup brain, isolating the cortex, and placing it in Hibernate E (Thermo Fisher, A1247601) on ice until cortices were harvested. Next, each cortex was minced into 1mm cubes and put in papain solution for digestion. Papain digest was stopped with 10% heat-inactivated fetal bovine serum (FBS), and each brain was triturated through a 10mL serological pipette to release neurons into the media. The media was then filtered through a 40 µM filter. Neurons were plated at a density of 750,000 cells per well on a poly-D-lysine coated 6-well plate. For 1 hour, neurons were cultured in a mix of DMEM: F12 with 10% FBS and pen/strep to facilitate adherence and early growth. After then neurons were cultured in a combination of Neurobala A with B-27 supplement, pen/strep, and ∼0.2X glutamax. Half media was changed on these neurons once per week. Neurons were cultured for 10 days, at which point they were harvested for experiments.

### Glycerol gradient analysis

Glycerol analysis on affinity-purified AUTS2 was conducted as previously described with certain modifications (Gao et al., 2012). Briefly, 12 ml nuclear extract (NE) from HEK293T-REx cells expressing FLAG-HA-AUTS2-S were incubated with 300 μl pre-washed FLAG M2 beads (Sigma, cat# A2220) and 3 ml Buffer A (10 mM Tris-HCl, pH 7.9, 1.5 mM MgCl_2_, 10 mM KCl, 0.5 mM DTT, 0.2 mM PMSF, 1 μg/ml Pepstatin A, 1 μg/ml Leupeptin, 1 μg/ml Aprotinin), in the presence of 0.02% NP-40. After an overnight rotation at 4 °C, the M2 beads were washed once with Buffer W (2/3 volume of Buffer C (20 mM Tris-HCl, pH 7.9, 25% glycerol, 420 mM NaCl, 1.5 mM MgCl_2_, 0.2 mM EDTA, 0.5 mM DTT, 0.2 mM PMSF, 1 μg/ml Pepstatin A, 1 μg/ml Leupeptin, 1 μg/ml Aprotinin), 5 times in 1/3 volume of Buffer A, NP-40 0.02%), then eluted with 600 μl FLAG peptides of 250 μg/ml in Buffer W by rotating at 4 °C for 1 hour. Subsequently, 500 μl M2 eluate was added to the top of a 12 ml 15 - 35 % glycerol gradient and centrifuged in an SW40Ti rotor (Beckman) at 40, 000 RPM at 4 °C for 22 hours. The resulting gradient was fractionated every 500 μl and then analyzed by immunoblotting.

### Immunoprecipitation

Immunoprecipitation (IP) experiments were performed as described previously (Gao et al., 2012) with modifications. NE from transfected HEK293T cells was incubated with 15 μl M2 beads at 4 °C overnight in a volume of 800 μl Buffer C, supplemented with 400 μl Buffer BN (20 mM Tris-HCl, pH 8.0, 100 mM KCl, 0.2 mM EDTA, 20% glycerol, 0.5 mg/ml BSA, 0.1% NP-40, 0.5 mM PMSF, 1 μg/ml Pepstatin A, 1 μg/ml Leupeptin, 1 μg/ml Aprotinin). Beads were then washed with Buffer BN 5 times and eluted with 50 μl FLAG peptides at 0.2 mg/ml at 4 °C for 1 hr. The eluates were mixed with SDS sample buffer and analyzed by SDS-PAGE, followed by immunoblotting.

### *In vivo* poly-ubiquitination assay

Examination of *in vivo* poly-ubiquitination of SMAD1 was performed as described previously (Abbas et al., 2008). Briefly, transfected cells with or without treatment were harvested directly in the buffer containing 1% SDS, 1mM EDTA, 2mM Na_3_VO_4_ in PBS. After 5 minutes of boiling, samples were passed through 26G needles 3 times and boiled for another 3 minutes. Next, the lysates were chilled and centrifuged, and the resulting supernatants were then mixed with equal volumes of Immunomix buffer (1% TX100, 1% SDS, 0.5% Deoxycholic acid, 1% BSA, 1mM EDTA, 2mM Na_3_VO_4_ in PBS) and subjected to immunoprecipitation with Anti-FLAG M2 beads, followed by SDS-PAGE and immunoblotting.

### Lentiviral infection

Lentivirus was generated in HEK293T cells through co-transfection of psPAX2, pMD136, and pLV-EF1a-IRES-Blast plasmid carrying genes of interest, using lipofectamine 2000 (Invitrogen, 11668019). Media containing lentivirus was collected, centrifuged at 4,000 g/4 °C for 15 minutes, and the resulting supernatant was concentrated on a spin column (EMD Millipore Corp, UFC910008). Concentrated lentivirus was aliquoted, fast-frozen in liquid nitrogen, and stored at - 80 °C. To infect mESCs with lentivirus, 300 μl condensed lentivirus was added to 6 ml ES media with 48 μg polybrene (EMD Millipore Corp, TR-1002-G). After 6 hours of incubation, media was removed and the cells were incubated overnight with a mixture of 10 ml media, 500 μl lentivirus, and 80 mg polybrene. The next day, infected mESCs were ready for further experiments.

### Alkaline phosphatase assay

AP staining was performed using a Stemgent AP staining kit II (ReproCELL, cat# 00-0055). Cells were fixed with fixation solution for 5 minutes at room temperature. After washing twice with PBS, the cells were incubated for 10 minutes in the dark with AP substrate solution. The reaction was terminated by removing the solution; the cells were then washed twice and covered with PBS for imaging.

### MTT proliferation assay

Cell proliferation was assayed according to the manufacturer manual (Vybrant® MTT Cell Proliferation Assay Kit, Invitrogen cat# M6494)). Briefly, cells were seeded in gelatin-coated 96-well plates with the same density in replicates. Absorbance at 540 nm wavelength was determined on a microplate reader (Bio-Rad).

### BMP treatment

Before treatment, cells were incubated in a serum-free starvation medium (1% BSA in DMEM) for 4 hours. BMP4 (R & R&D Systems, cat#314-BP-010) was then added with a 5 ng/ml final concentration. Cells were collected at different time points after BMP4 treatment and subjected to immunoblotting. The image was quantified by densitometry using Image J. The mean grey value of a selected region that contained an individual band was measured and subtracted by the background.

### Statistics

Statistical analyses were performed using unpaired, two-tailed Student’s t-test where applicable for comparison between two groups. All data are presented as mean ± standard deviation of the mean. The sample size for each experiment is included in the figure legends. Statistical significance was defined as *p < 0.05, **p < 0.01, ***p < 0.001; ns, not significant.

