## Supplemental Infomation for "AUTS2 controls neuronal lineage choice through a novel PRC1-independent complex and BMP inhibition"

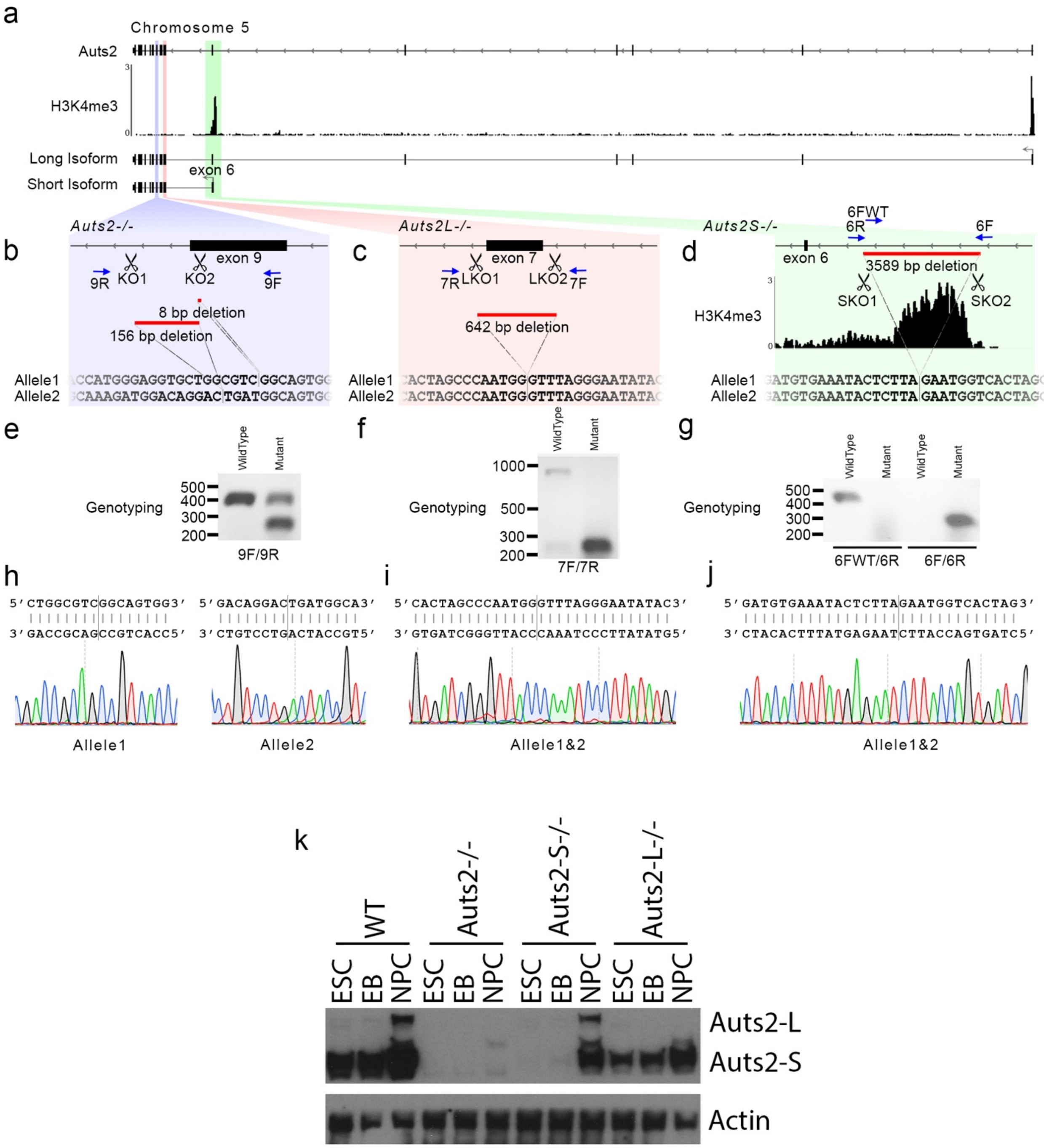
a
Chromosome 5
Auts2
H3K4me3
Long Isoform
Short Isoform
exon 6
b
Auts2-/-
exon 9
KO1
KO2
9R
9F
8 bp deletion
156 bp deletion
Allele1
Allele2
c
Auts2L-/-
exon 7
LKO1
LKO2
7R
7F
642 bp deletion
d
Auts2S-/-
6FWT
6R
6F
exon 6
3589 bp deletion
SKO1
SKO2
e
Genotyping
WildType
Mutant
9F/9R
f
7F/7R
g
6FWT/6R
6F/6R
h
Allele1
Allele2
i
Allele1&2
j
Allele1&2
k
WT
Auts2-/-
Auts2-S-/-
Auts2-L-/-
ESC
EB
NPC
Auts2-L
Auts2-S
Actin

**Figure S1. Design, generation and validation of mouse ESC lines with Auts2 deletion.** ***a***, Schematic representation of the genetic locus of mouse *Auts2*. H3K4me3 peaks, indicating the two transcription start sites, were obtained from previous ChIP-seq analysis [6]. Two major transcripts, one referred to as “long” and the other referred to as “short” isoforms are shown at the bottom. ***b-d***, Targeting strategy and final products of CRISPR/Cas9-mediated gene editing of *Auts2* locus. In each cell line, a pair of sgRNAs (labeled by scissors) were used and the deleted regions are indicated by red bars. Blue arrows indicate genotyping primers. Post-editing sequences are shown for both alleles. The two alleles in the *Auts2-/-* line were edited differently; one has an 8 bp frameshift deletion and the other a 156 bp deletion that spans the splicing junction (***b***). Identical editing events are present in both *Auts2L-/-* (***c***) and *Auts2S-/-* (***d***) lines. ***e-g***, Genotyping PCR results for edited lines. ***h-j***, Sanger sequencing results for edited lines. ***k***, Immunoblotting to confirm the disruption of specific isoforms of *Auts2* in edited lines.

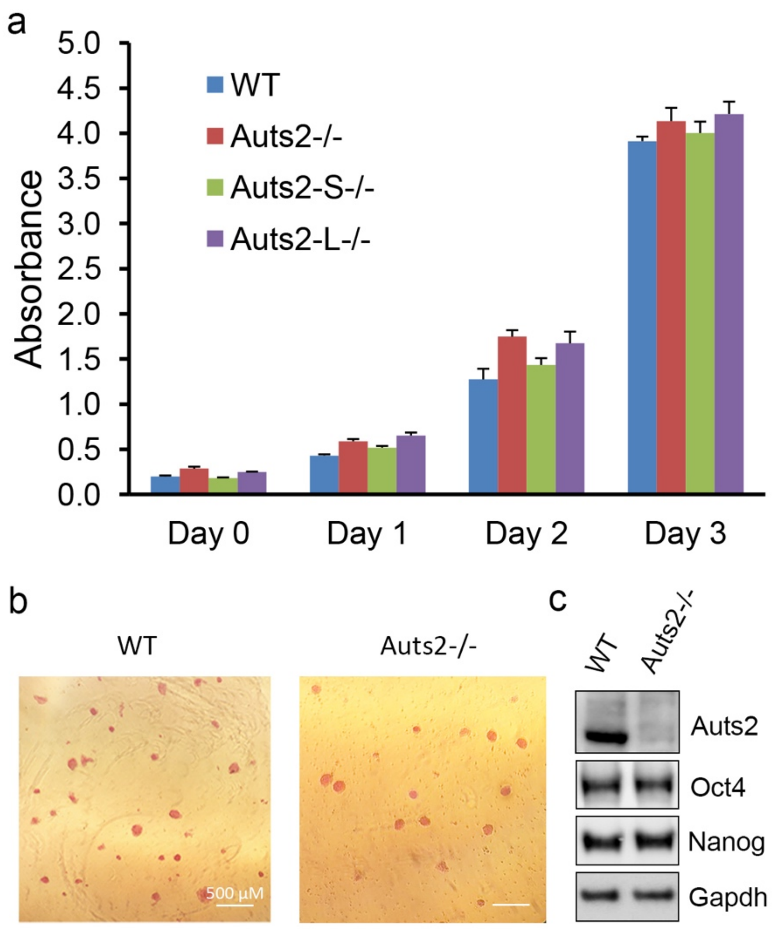


**Figure S2. Loss of Auts2 does not affect ESC self-renewal and proliferation.** ***a***, Alkaline phosphatase activity staining in WT and *Auts2*-/- ESCs. ***b***, Immunoblotting shows no obvious difference in the protein level of Oct4 and Nanog, two pluripotent markers, between WT and *Auts2*-/- ESCs. ***c***, MTT cell proliferation assay to measure the growth rate of ESCs at various days. All mean values and standard deviations were calculated from four independent measurements. No significant differences were found among ESCs of all genotypes.

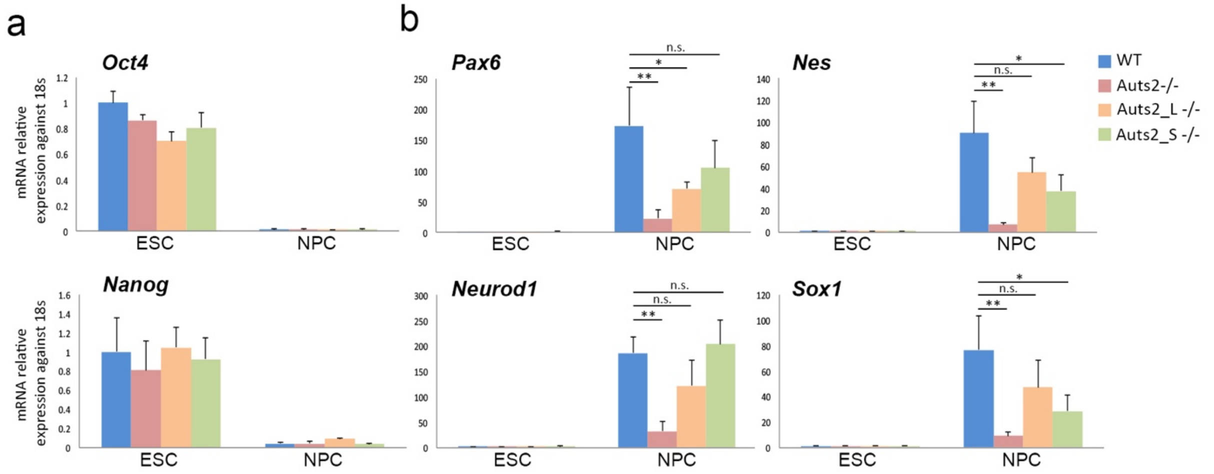


**Figure S3. Expression of pluripotency and NPC markers in ESCs with Auts2 deletion**. ***a***, Quantitative RT-PCR analysis of expression pluripotency markers (*Oct4* and *Nanog*) in WT, *Auts2*-/-, *Auts2L*-/- and *Auts2S*-/- ESCs and NPCs. ***b***, Quantitative RT-PCR analysis of NPC markers (*Pax6*, *Nes*, *Neurod1* and *Sox1*). Expression levels are normalized relative to those in WT ESCs. All mean values and standard deviations were calculated from three independent measurements. * $P<0.05$, ** $P<0.01$, n.s., not significant, by two-sided t-test.

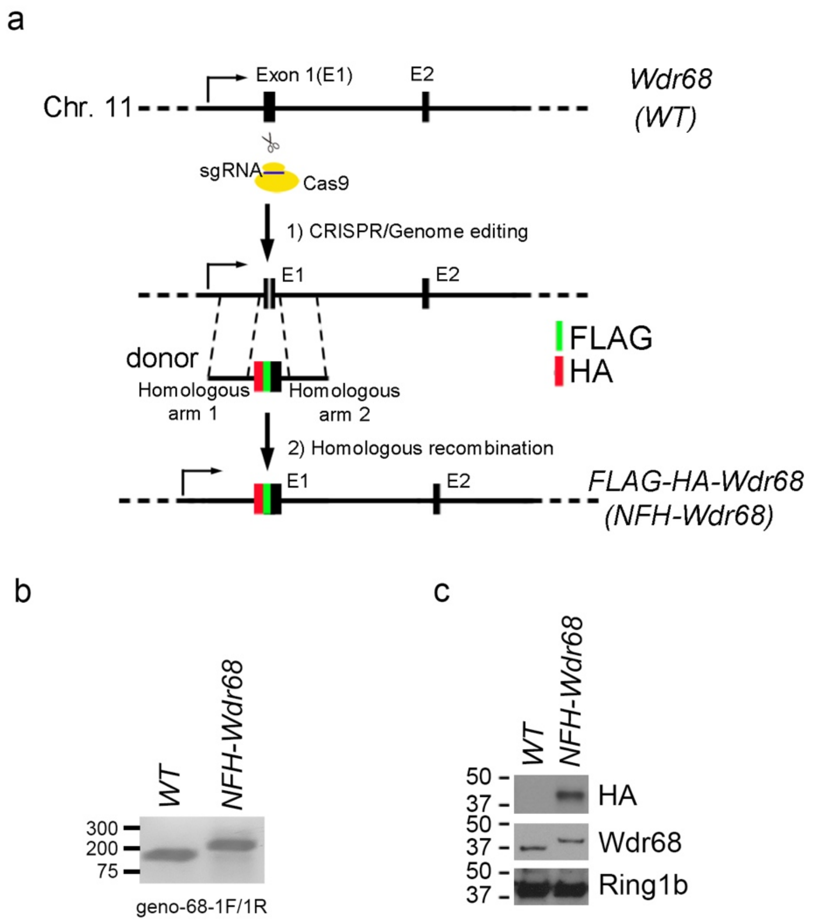


**Figure S4. Generation of an ESC line that expresses FLAG-HA-Wdr68.** ***a***, Schematic for the CRISPR/Cas9-mediated knock-in of FLAG-HA-Wdr68 in E14 cells. ***b***, Agarose gel picture shows the PCR products of genomic DNA isolated from WT and FLAG-HA-Wdr68 cells. ***c***, Immunoblotting detected the expression of FLAG-HA-Wdr68 in edited cells.

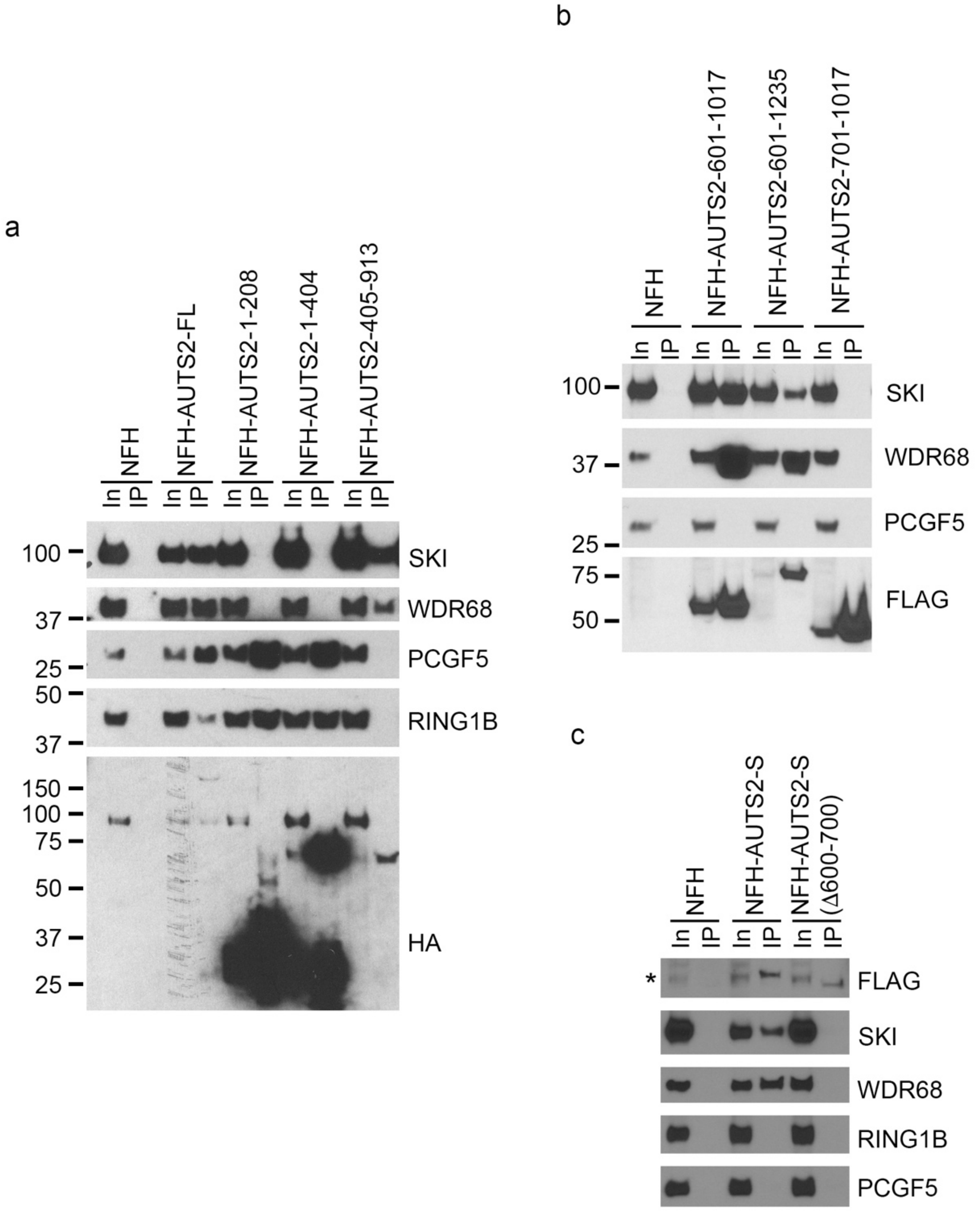


**Figure S5. Domain mapping of AUTS2 for interaction with various partner proteins.** ***a***, ***b*** and ***c***, HEK293T cells were plasmids expressing various lengths of FLAG and HA tagged AUTS2 as indicated. IP was performed using M2 beads. Bound proteins were resolved on SDS-PAGE and detected by Western blotting for the indicated antigens.

### Table S1 Plasmids

| | Plasmid Name | Source | Vector | Enzyme sites | Tags |
|---|---|---|---|---|---|
| 1 | pINTO-NFH-AUTS2-L | Gao et al., 2014 | pINTO-NFH | KpnI/EcoRI | N-FLAG-HA |
| 2 | pINTO-NFH-AUTS2-S | This study | pINTO-NFH | | N-FLAG-HA |
| 3 | pINTO-NFH-AUTS2-8-208 | This study | pINTO-NFH | KpnI/XhoI | N-FLAG-HA |
| 4 | pINTO-NFH-AUTS2-8-404 | This study | pINTO-NFH | KpnI/XhoI | N-FLAG-HA |
| 5 | pINTO-NFH-AUTS2-405-913 | This study | pINTO-NFH | KpnI/XhoI | N-FLAG-HA |
| 6 | pINTO-NFH-AUTS2-601-1017 | This study | pINTO-NFH | | N-FLAG-HA |
| 7 | pINTO-NFH-AUTS2-601-1235 | This study | pINTO-NFH | | N-FLAG-HA |
| 8 | pINTO-NFH-AUTS2-701-1017 | This study | pINTO-NFH | | N-FLAG-HA |
| 9 | pINTO-NFH-AUTS2-701-1235 | This study | pINTO-NFH | | N-FLAG-HA |
| 10 | HA-SKI | Addgene, #10910 | pCI-Neo | XhoI/HpaI | N-HA |
| 11 | HA-DDB1 | Addgene, #19909 | pCDNA3-HA2 | | N-HA |
| 12 | HA-ub | Iwahara et al., 2012 | | | HA |
| 13 | FLAG-SMAD1 | Addgene, #11735 | pCMV5B | ClaI/BamHI | N-FLAG |
| 14 | DN-CUL4B | Addgene, #15822 | pCDNA3.1(+) | BamHI/ApaI | C-FLAG |
| 15 | pLV-EF1a-IRES-Blast | Addgene, #85133 | pLV-EF1a-IRES-Blast | BamHI/EcoRI | |
| 16 | pLV-hAUTS2-S | This study | pLV-EF1a-IRES-Blast | BamHI/EcoRI | |
| 17 | pLV-hAUTS2-Δ | This study | pLV-EF1a-IRES-Blast | BamHI/EcoRI | |

**Table S2 Antibodies used in this study**

| Antibody | Suppliers | Catalog No. | Lot No. | Applications |
|---|---|---|---|---|
| AUTS2 | Gao et al., 2014 | | | WB, IF |
| AUTS2 | Sigma | HPA000390 | | WB |
| WDR68 | Sigma | HPA022948 | A76005 | WB |
| SKI | Santa Cruz | sc33693 | | WB |
| SMAD1 | Cell Signaling | 6944 | lot: 5 | WB |
| SMAD2 | Cell Signaling | 5339 | lot: 4 | WB |
| SMAD4 | Cell Signaling | 38454 | lot: 1 | WB |
| SMAD5 | Cell Signaling | 12534 | lot: 2 | WB |
| pSMAD2 | Cell Signaling | 3108 | lot: 8 | WB |
| pSMAD1/5/9 | Cell Signaling | 13820 | lot: 3 | WB |
| RING1B | BETHYL | A302-869A | A302-869A-1 | WB |
| PCGF3/5 | Abcam | ab201510 | | WB |
| PCGF5 | ABCAM | ab201511 | | WB |
| DDB1 | Santa Cruz | sc376860 | A2318 | WB |
| NANOG | Invitrogen | PA1-41577 | | WB |
| OCT4 | Santa Cruz | sc365509 | L1817 | WB |
| PAX6 | DSHB | Pax6-s | | WB |
| NESTIN | BD | 611658 | | WB, IF |
| HA | Covance | MMS-101P | | WB |
| HA | Abcam | ab9110 | GR98618-3 | WB |
| HA beads | Sigma | A2095 | | IP |
| FLAG | Sigma | F3165 | | WB |
| FLAG beads | Sigma | A2220 | | IP |
| β-TUBULIN | Abcam | ab6046 | | WB |
| GAPDH | Thermo Fisher Scientific | MA5-15738 | GA1R | WB |
| ACTIN | Abcam | ab8277 | | WB |
| H3 | Abcam | ab1791 | | WB |

**Table S3 List of primers for qPCR**

**For RT-PCR**

| Name | Sequence (F/R) |
|---|---|
| *mOct4* | 5'-AGATCACTCACATCGCCAATCA-3' / 5'-CGCCGGTTACAGAACCATACTC-3' |
| *mSox9* | 5'-AGCGAACGCACATCAAGA-3' / 5'-CTGTAGTGAGGAAGGTTGAAGG-3' |
| *mNanog* | 5'-AGGCTTTGGAGACAGTGAGGTG-3' / 5'-TGGGTAAGGGTGTTCAAGCACT-3' |
| *mNes* | 5'-AGTGCCCAGTTCTAGTGGTGTCC-3' / 5'-CCTCTAAAATAGAGTGGTGAGGGTTG-3' |
| *mTbx2* | 5'-ATGTACATCCACCCGGACAG-3' / 5'-GACAGCGATGAAGTCGGTCT-3' |
| *mDpysl2* | 5'-CAGAATGGTGATTCCCGGAGG-3' / 5'-CAGCCAATAGGCTCGTCCC-3' |
| *mSnai1* | 5'-CCGATGAGGACAGTGGCAAA-3' / 5'-CCCAGGCTGAGGTACTCCTT-3' |
| *mNeurod1* | 5'-CGAGTCATGAGTGCCCAGCTTA-3' / 5'-CCGGGAATAGTGAAACTGACGTG-3' |
| *mId2* | 5'-CTACTCCAAGCTCAAGGAACTG-3' / 5'-GATCTGCAGGTCCAAGATGTAA-3' |
| *mPax6* | 5'-CTTGGGAAATCCGAGACAGA-3' / 5'-CTAGCCAGGTTGCGAAGAAC-3' |
| *m18SrRNA* | 5'-GCAATTATTCCCCATGAACG-3' / 5'-GGCCTCACTAAACCATCCAA-3' |
| *mTubb2a* | 5'-GGAGGTGATAAGCGATGAGCATG-3' / 5'-GGCTCCAGGTCCACTAGGATG-3' |
| *mHoxa6* | 5'-GTGACCCTACTGCCATCTTAC-3' / 5'-GAATAATCACCGCAGGACTCT-3' |
| *mWwc2* | 5'-TCATCTGGGAGCAGTCTAGGG-3' / 5'-TGATAGTCTGTGTCCATCTGGTC-3' |
| *mHand1* | 5'-TCGCTACACTTCCTACCTAGAG-3' / 5'-GAAGGGAAAGGAAGGGAAAGAT-3' |
| *mTubb3* | 5'-TGAGGCCTCCTCTCACAAGT-3' / 5'-GGCCTGAATAGGTGTCCAAA-3' |
| *mGata6* | 5'-TGCAGGATTGCATCATGACAGA-3' / 5'-TGACCTCAGATCAGCCACGTT-3' |
| *mVglut1* | 5'-TGCTACCTCACAGGAGAATGGA-3' / 5'-GCGCACCTTCTTGCACAAAT-3' |
| *mGap43* | 5'-TGTGCCTGCTGCTGTCACTGAT-3' / 5'-AGGTTTGGCTTCGTCTACAGCG-3' |
| *mGata4* | 5'-TTCCTCTCCCAGGAACATCAAA-3' / 5-GCTGCACAACTGGGCTCTACTT-3' |
| *mSox1* | 5'-GGCCGAGTGGAAGGTCAT-3' / 5'-ACTTGTAATCCGGGTGTTCCT-3' |
| *mNkx2-5* | 5'-GATGGGAAAGCTCCCACTATG-3' / 5'-GACACCAGGCTACGTCAATAAA-3' |
| *mT(Brachyury)* | 5'-TGCACATTACACACCACTGACG-3' / 5'-AGAACCAGAAGACGAGGACGT-3' |
| *mMap2* | 5'-AAAGGCCCGCGTAGATCAC-3' / 5'-GGGATTCGAGCAGGTTGATG-3' |

**For geno typing**

| Name | Sequence |
|---|---|
| *mWdr68KI-1F* | 5'-CAGCCCGCTGCCTCTCTGG-3' |
| *mWdr68KI-1R* | 5'-TTCCACGAAGCTGCCCAGCG-3' |
| *mAuts2KO-7F* | 5'-TTTTATAGATTTTTCTTTATCCACATGT-3' |
| *mAuts2KO-7R* | 5'-GGTGTGTGCCATCATTCCTGGG-3' |
| *mAuts2KO-9F* | 5'- GGAACTGAACACGCGTTTCCTG-3' |
| *mAuts2KO-9R* | 5'-CACAGTGCCAGCAATGGCCATC-3' |
| *mAuts2KO-6F* | 5'-GCTGGTGTGAATGTATACACAAGT-3' |
| *mAuts2KO-6R* | 5'-TGGCACCAAAGGTGTTACTATTCC-3' |
| *mAuts2KO-6FWT* | 5'-CCTAATCCAGCTCTGGCTCCCA-3' |
| *mSKI-F* | 5'-AAAGATGTGCGCCCAGTGCGCA-3' |
| *mSKI-R* | 5'-CAGGATGCCCATGACTTTGAGGA-3' |
